# Large MAF Transcription Factors Reawaken Evolutionarily Dormant Fast-Glycolytic Type IIb Myofibers in Human Skeletal Muscle

**DOI:** 10.1101/2024.12.09.627660

**Authors:** Shunya Sadaki, Ryosuke Tsuji, Takuto Hayashi, Masato Watanabe, Ryoto Iwai, Gu Wenchao, Ekaterina A. Semenova, Rinat I. Sultanov, Andrey V. Zhelankin, Edward V. Generozov, Ildus I. Ahmetov, Iori Sakakibara, Koichi Ojima, Hidetoshi Sakurai, Masafumi Muratani, Takashi Kudo, Satoru Takahashi, Ryo Fujita

## Abstract

Small mammals rely on type IIb myofibers, expressing the fastest myosin IIb (encoded by *MYH4*), for rapid muscle contraction. In contrast, larger mammals, including humans, show reduced or absent *MYH4* expression and type IIb myofibers, favoring slower-contracting myofibers. The evolutionary mechanisms underlying this shift remain unclear. Here, we identify large MAF transcription factors (MAFA, MAFB, MAF) as key regulators of *MYH4* expression in large mammals, including human and bovine. Overexpression of large MAFs induces MYH4 expression and enhances glycolytic capacity in human myotubes, supported by RNA-seq and metabolic flux analyses. RNA-seq of human muscle biopsies reveals a positive correlation between *MAFA*, *MAF*, and *MYH4* expression, with these genes elevated in power-trained individuals. These findings reveal a conserved mechanism across mammals, showing that large MAFs can induce type IIb myofibers even in humans, with potential applications for enhancing athletic performance and addressing age-related muscle weakness associated with the loss of fast-twitch myofibers.

## Introduction

Skeletal muscles are composed of myofibers with distinct contractile, metabolic, and fatigue-resistant properties, as well as differing vulnerabilities in pathophysiological conditions (*1*, *2*). Myofibers are classified as slow-twitch (type I) and fast-twitch (type II). Type I myofibers are rich in mitochondria, exhibit high oxidative metabolism, and express slow isoforms of sarcomeric proteins, such as myosin heavy chain 7 (*Myh7*). Type II myofibers are further divided into types IIa, IIx, and IIb based on the expression of the Myh isoforms *Myh2*, *Myh1*, and *Myh4*, respectively. Each myosin isoform exhibits different ATPase activities that correlate with muscle contraction speed. Fast myofibers that express *Myh4* contract at greater speeds than those expressing *Myh1, Myh2*, and *Myh7*, consequently generating up to ten times more power than slow myofibers (*3*, *4*). Additionally, type IIb myofibers have an underdeveloped mitochondrial network and high concentrations of glycolytic enzymes, thereby supporting active glycolysis for ATP production.

Skeletal muscle is the most abundant tissue in adult humans, accounting for approximately 40% of the total body mass. Consequently, the proportion of myofiber types affects muscle performance and whole-body metabolism. In addition, it affects susceptibility to conditions such as diabetes (*5*, *6*) and the progression of muscular dystrophy (*7*, *8*). Myofiber composition can be remodeled in response to various pathophysiological conditions, including exercise, disuse, diabetes, space travel, and aging. During disuse-induced muscle atrophy, as observed in space travel or immobilization, a slow-to-fast type IIb conversion is exhibited in the mouse soleus muscle, which primarily consists of type I and type IIa myofibers (*9*, *10*). Conversely, aerobic exercise can activate slow muscle gene programs, including PGC1α, in rodents (*11–13*). Myofiber types substantially change with aging. Aging induces a type IIb-to-IIx shift in fast muscles and a type IIa-to-I shift in slow muscles in rodent skeletal muscle. This results in an overall fast-to-slow myofiber type transition (*14*, *15*). Human type II myofibers are more prone to myonuclear apoptosis and atrophy with aging, leading to a fast-to-slow transition in sarcopenic muscle (*16*, *17*). This may contribute to decreased locomotor activity in the elderly. However, the mechanistic role of myofiber type shifts in human skeletal muscle pathogenesis, including aging, remains largely unknown.

The composition of Myh isoforms is highly diverse across animal species. Previous studies characterizing Myh isoforms across species have revealed a notable distinction between large and small mammals regarding *Myh4* expression (*16*, *18–20*). Small animals such as rodents contain the full spectrum of myofiber types, including type I, IIa, IIx, and IIb myofibers. However, the muscles of larger animals, including humans, typically lack type IIb myofibers and low expression or complete absence of the *MYH4* (*20*, *21*). Instead, human skeletal muscles express only type I, IIa, and IIx myofibers, along with their corresponding MYH genes *MYH7*, *MYH2*, and *MYH1*, respectively. A small subset of specialized muscles, such as extraocular muscles in humans, exhibit detectable levels of *MYH4* mRNA, but not the protein (*22*, *23*). As large animals retain the *MYH4* gene, we hypothesized that they can develop type IIb myofibers by activating *MYH4* expression. Type IIb myofibers generally enable rapid contraction, are larger than other myofibers, and generate a strong force with high glycolytic activity (*17*, *24*). Therefore, understanding why large animals abandoned *MYH4* expression during evolution could provide insights into muscle function and adaptation across species, particularly as it relates to physiological and metabolic demands unique to large animals. Furthermore, elucidating the mechanisms underlying the loss of type IIb myofibers may uncover strategies to re-induce these powerful, rapid-contracting fibers in humans. These advances could facilitate the enhancement of human muscle function and performance, potentially extending physical capabilities in both clinical and athletic settings.

The adult fast myosin (fMyh) genes *Myh2*, *Myh1*, *Myh4* are clustered on chromosomes 11 and 17 in mice and humans, respectively. Dos Santos et al. (*25*) identified a 42-kb fMyh super-enhancer (fMyh-SE) region that interacts with the fMyh promoter through 3D chromatin looping. However, the mechanisms by which transcription factors regulate the fine-tuned expression of each fMyh gene within myofibers via this enhancer region have not been fully elucidated. Our previous study was the first to successfully identify the large Maf transcription factor family (hereafter referred to as Large Mafs in mice and large MAFs in humans) as a principal regulator of type IIb myofibers (*26*). Although four large Mafs— *Mafa*, *Mafb*, *Maf*, and *Nrl*—have been identified in mammals, only *Mafa*, *Mafb*, and *Maf* are expressed in the skeletal muscle of mice and humans (*26*). Furthermore, single deletion of *Mafa*, *Mafb*, or *Maf* in skeletal muscle does not affect the proportion of myofiber types. However, deletion of all three large Mafs in skeletal muscle leads to an almost complete loss of type IIb myofibers. This increases endurance capacity and reduces muscle force in mice (*26*). Conversely, overexpression of each large Maf induced type IIb myofibers in the soleus muscle by binding to the Maf recognition element located −270bp upstream of the transcription start site of *Myh4* in adult mice. These findings indicate that large Mafs are a potent and specific regulator of type IIb myofiber determination in mice (*26*). However, their role in the skeletal muscle of larger animals, including humans, remains completely unknown. Moreover, the induction of fast-glycolytic type IIb myofibers—long lost in human skeletal muscle—through the manipulation of large MAFs remains a major scientific challenge.

Here, we report for the first time that *MYH4*, the corresponding gene of type IIb fibers, which is typically not expressed at the mRNA or protein level in humans, is robustly and specifically induced, by 100- to 1000-fold, in human and bovine skeletal muscle cells through overexpression of large MAFs. Importantly, we demonstrate that MYH4 protein, previously undetectable in human muscle, was translated in human myotubes and detected by LC-MS/MS analysis. Furthermore, overexpression of large MAFs profoundly alters the expression of genes associated with glycolytic metabolism, thereby enhancing glycolytic activity in human skeletal muscle cells. Analysis of human muscle biopsy samples also revealed a positive correlation between *MAFA* and *MAF* expression and the proportion of fast-twitch myofibers in human skeletal muscle. Notably, *MAFA* and *MAF* expression was also positively correlated with *MYH4* expression in human skeletal muscles, with these genes elevated in power-trained individuals. Our findings could enable the enhancement of human muscular strength and functional capacity by inducing Type IIb-like properties similar to those found in the muscles of small animals such as mice and rabbits, thereby unlocking new possibilities for human muscle performance.

## Results

### Positive Correlation between the Expression of large MAFs and MYH4 Expression Across Animal Species

To investigate why large mammals lack type IIb myofibers and the corresponding *MYH4* gene, we analyzed the expression levels of *MYH4* (human), *Myh4* (mouse), and their orthologs in other species, as well as those of large MAF genes (*MAFA*, *MAFB*, and *MAF*), across various animal species using the Bgee database (*27*). The vertical axis represents the expression score of *MYH4*/*Myh4*, while the horizontal axis depicts the combined mRNA expression scores of *MAFA*, *MAFB*, and *MAF*, revealing a positive correlation between large MAF mRNA expression and *MYH4*/*Myh4* mRNA expression across species (Fig. 1A). Notably, small animals such as mice, rats, opossums, and frogs exhibited elevated expression of large Mafs. Conversely, large mammals, including horses, bovines, and humans, showed reduced total large MAF expression scores. These data suggest that the expression levels of large MAFs determine the presence of type IIb myofibers across diverse animal species.

**Fig. 1.**
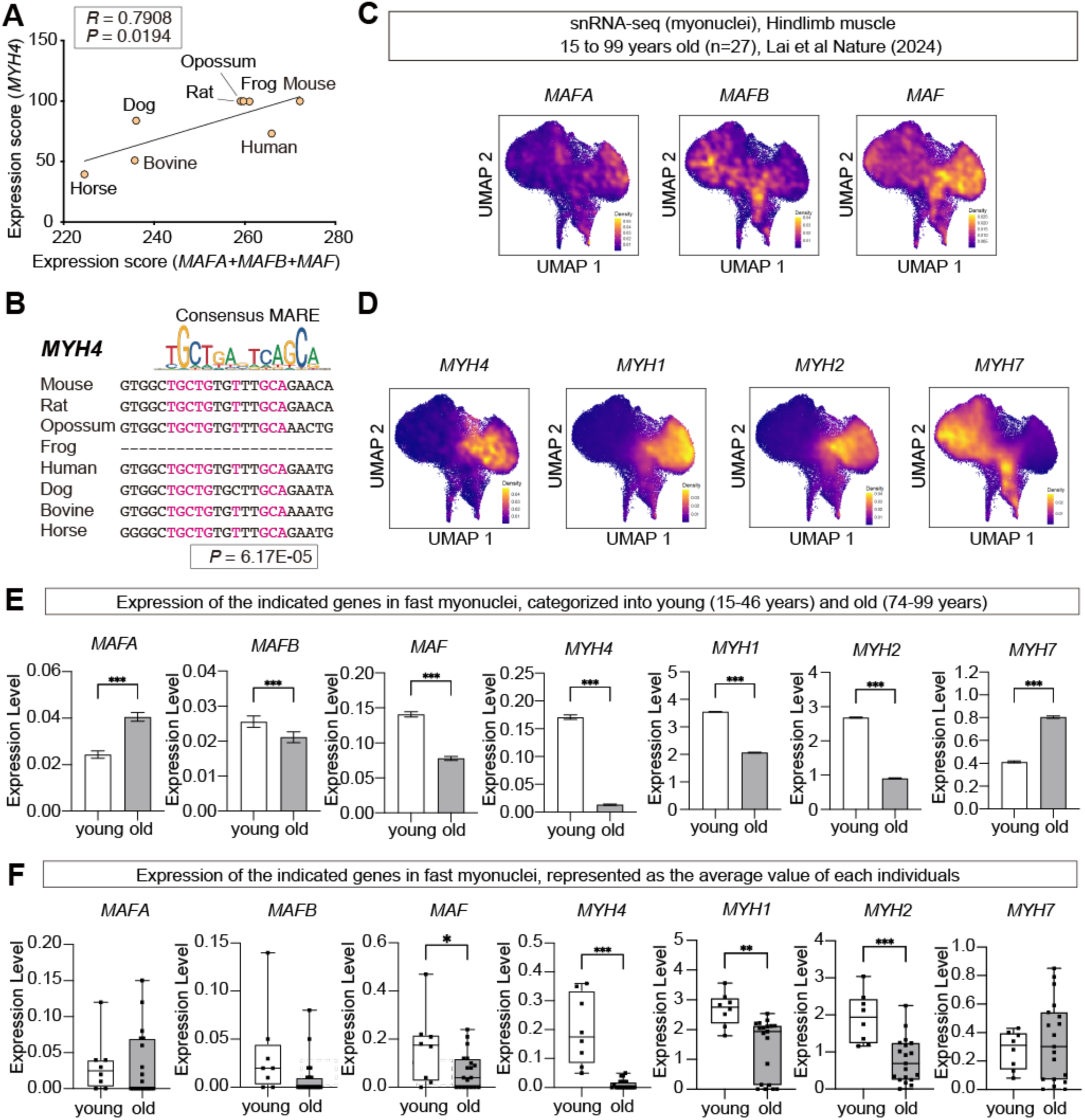
MYH4 expression fluctuates in response to large MAFs expression. (**A**) Correlation between the expression of large MAFs (*MAFA*, *MAFB*, and *MAF*) and *MYH4* across multiple animal species. (**B**) Presence of MARE-like sequences in the promoter region of *Myh4* across multiple animal species. (**C**) Visualization of *MAFA*, *MAFB*, and *MAF* expression using UMAP, as derived from single-nucleus RNA sequencing (snRNA-seq) of human hindlimb muscle samples from individuals aged 15–99 years (n = 27). Data sourced from Lai et al., *Nature* (2024). (**D**) Visualization of *MYH4*, *MYH1*, *MYH2*, and *MYH7* expression using UMAP, as derived from snRNA-seq of human hindlimb muscle samples from individuals aged 15–99 years (n = 27). Data sourced from Lai et al., *Nature* (2024). (**E**) Expression levels of *MAFA*, *MAFB*, *MAF*, *MYH4*, *MYH1*, *MYH2*, and *MYH7* in all myonuclei, categorized by age group: young (15–46 years) and old (74–99 years). (**F**) Average expression levels of *MAFA*, *MAFB*, *MAF*, *MYH4*, *MYH1*, *MYH2*, and *MYH7* in myonuclei, represented as the mean value for each individual in the respective age groups (young: 15–46 years, n = 8; old: 74–99 years, n = 19). P values were calculated using a two-sided Pearson correlation test (**A**) and an unpaired two-tailed *t*-test (**E**, **F**); *P < 0.05, **P < 0.01, ***P < 0.001, ****P < 0.0001, R; Pearson correlation coefficient.

Large Mafs directly induce type IIb myofibers via a MARE site within the *Myh4* promoter region in mice (*26*). To further explore this regulatory mechanism, we examined the MARE sequences in the *MYH4* promoter across several animal species to assess the potential of large MAFs to regulate *MYH4* expression in large mammals, including humans (Fig. 1B). With the exception of the amphibian frog, we found that the MARE sequences in the *MYH4* promoter were highly conserved, even among large mammals (Fig. 1B). These findings suggest that the upregulation of large MAFs levels could induce *MYH4* expression through the MARE in large animals, including humans.

### *MAFA* and *MAF* Enriched in Fast Myonuclei in Human Skeletal Muscle Decline with Aging

We next analyzed single-nucleus RNA sequencing (snRNA-seq) data of human skeletal muscle (*16*) to evaluate the role of large MAFs. Although *MYH4* expression was minimal in human skeletal muscle, *MAFA* and *MAF* were enriched in fast myonuclei expressing *MYH2*, *MYH1*, and *MYH4* (Fig. 1C, D). While *MAFB* was also expressed in myonuclei, its expression was broadly distributed across myonuclei expressing both fast and slow *MYH* genes (Fig. 1C, D).

To examine the impact of aging on their expression, we compared snRNA-seq data from young individuals (15–46 years old) with those from elderly individuals (74–99 years old) (Fig. 1E, F). Analysis of *MAFA*, *MAFB*, *MAF* expression in fast myonuclei revealed a significant downregulation in *MAFB* and *MAF* expression in aged myonuclei, whereas *MAFA* expression was upregulated (Fig. 1E). Fast *MYH*, such as *MYH1*, *MYH2*, and *MYH4*, showed a substantial decline with aging (Fig. 1E). However, *MYH4* expression remained extremely low in both young and old myonuclei. In contrast, *MYH7* expression was upregulated in aged myonuclei compared to young myonuclei. We further analyzed the average expression levels of *MAFA*, *MAFB*, *MAF,* and fast/slow *MYH* for each individual in both cohorts (Fig. 1F) using the same snRNA-seq dataset (*16*). The expression of fast MYH genes (*MYH4*, *MYH1*, *MYH2*) was significantly reduced in older individuals, concomitant with decreased *MAF* expression (Fig. 1F). These results suggest that large MAFs regulate fast myofiber gene expression in human skeletal muscle and highlight their physiological relevance in human aging.

### Large MAFs Specifically Induce MYH4 Expression in Human Skeletal Muscle Cells

To investigate whether large MAFs play a role in inducing type IIb myofibers in humans, we overexpressed *MAFA*, *MAFB*, or *MAF* using adenoviral vectors in human skeletal muscle cells derived from human iPS cells established previously (*28*, *29*). Sorted induced muscle stem cells (iMuSCs) derived from iPS cells were differentiated for 2 days in differentiation medium (DM) before transduction with adenoviral particles expressing *MAFA*, *MAFB*, or *MAF* driven by the CMV promoter (Fig. 2A). An adenoviral vector expressing *mCherry* alone was used as control (CON) (Fig. 2A) No abnormalities were observed in myotubes overexpressing *MAFA*, *MAFB*, or *MAF* compared to the control myotubes following viral transduction (Fig. 2B). The infection efficiency, determined based on the percentage of mCherry-positive myotubes, was comparable across all groups (>90%) (Fig. 2B-C). Myotube width was unaffected by large MAF overexpression (Fig. 2D). Furthermore, *MAFA*, *MAFB*, and *MAF* were successfully overexpressed in human myotubes (Fig. 2E).

**Fig. 2.**
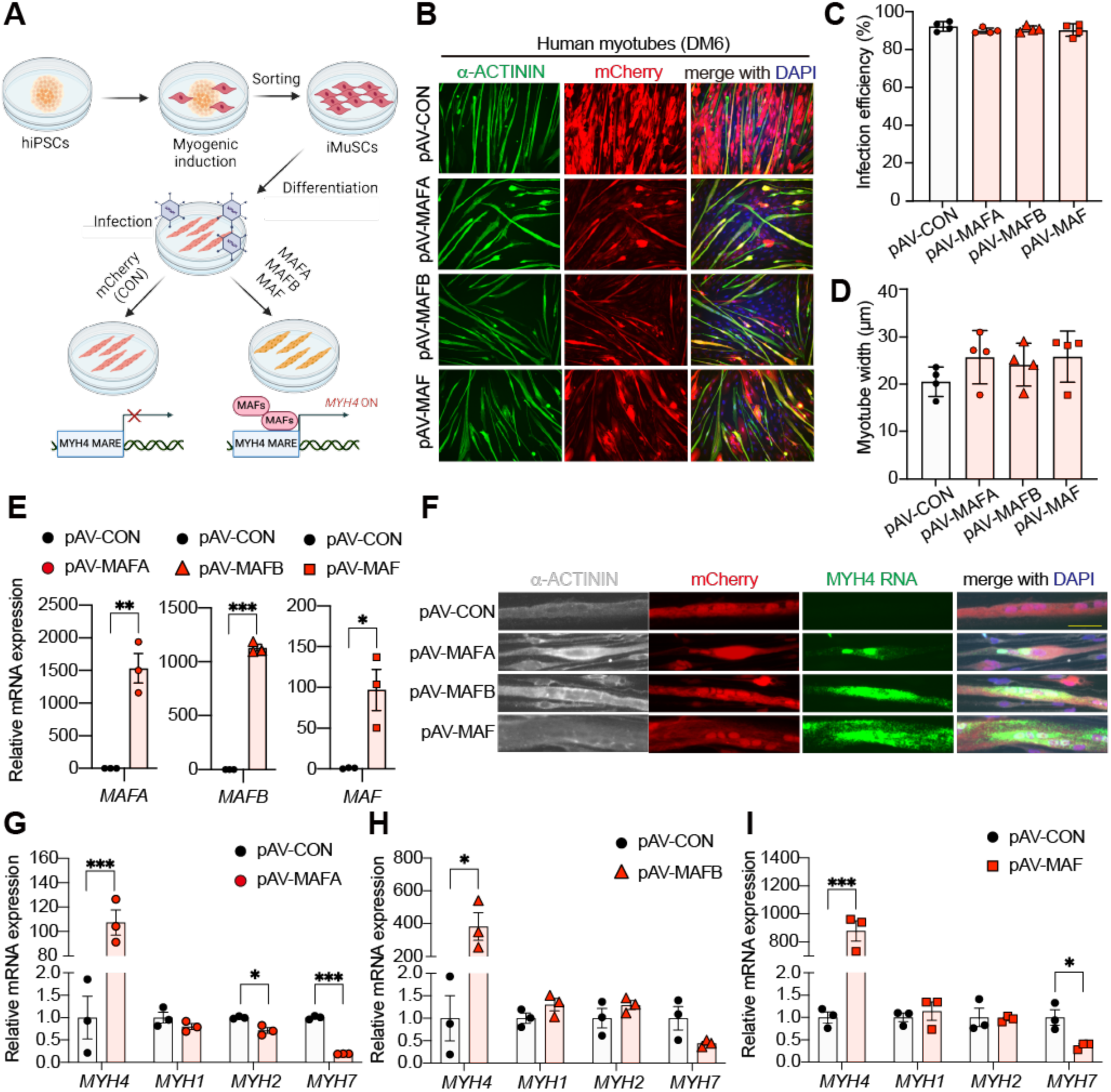
Large MAFs can specifically induce MYH4 mRNA expression in human iPSC-derived myotubes. (**A**) Schematic diagram illustrating adenovirus-mediated overexpression of MAFA, MAFB, and MAF in human iPSC-derived myotubes. After myogenic induction, iMuSCs were purified using flow cytometry and cultured in growth medium (GM) for expansion. After two days of exposure to differentiation medium (DM), human myotubes were treated with adenoviral particles expressing MAFA_mCherry (pAV-MAFA), MAFB_mCherry (pAV-MAFB), or MAF_mCherry (pAV-MAF). mCherry-only was used as the control (pAV-CON). Created in BioRender. Ryo, F. (2025) https://BioRender.com/r49s132. (**B**) Immunostaining of human myotubes transduced with pAV-CON, pAV-MAFA, pAV-MAFB, or pAV-MAF using antibodies against alpha-actinin (α-ACTININ, green) and mCherry (red). Nuclei were stained with DAPI (blue). (**C**) Quantification of infection efficiency for each adenoviral vector. The proportion of mCherry-positive cells within α-ACTININ-positive myotubes was calculated. In total, 91– 105 myotubes were analyzed per experiment (n = 4). (**D**) Quantification of myotube width shown in panel (**B**). In total, 157–283 myotubes were analyzed per experiment (n = 4). (**E**) Relative mRNA expression levels of MAFA, MAFB, and MAF, as determined using RT-qPCR in human myotubes 4 days after adenovirus transduction (n = 3/group). (**F**) Detection of MYH4 mRNA in human myotubes transduced with pAV-CON, pAV-MAFA, pAV-MAFB, or pAV-MAF using RNAscope. Myotubes were visualized with an antibody against α-ACTININ (white) and mCherry (red) after hybridization. Nuclei were stained with DAPI (blue). Scale bar: 50 μm. (**G-I**) Relative mRNA expression levels of myosin heavy chain genes (*MYH1*, *MYH2*, *MYH4*, and *MYH7*), as determined using RT-qPCR in human myotubes 4 days after transduction with adenoviral vectors expressing MAFA (**G**), MAFB (**H**), or MAF (**I**). mCherry-only was used as the control (pAV-CON, n = 3/group). P values were calculated using Tukey’s test (**C**, **D**) and Student’s *t*-test (**E**, **G-I**); *P < 0.05, **P < 0.01, ***P < 0.001.

We further performed RNAscope fluorescent in situ hybridization to determine whether large MAFs induce *MYH4* expression. *MYH4* RNA puncta were barely detectable in control human myotubes overexpressing mCherry (Fig. 2F). However, *MYH4* puncta were clearly accumulated in all large MAF-overexpressing myotubes (Fig. 2F). We used qPCR analysis to further quantify the expression levels of *MYH4* and other *MYH* genes. Although *MYH4* expression in control myotubes was markedly low, it was elevated by approximately 100- to 1000-fold in large MAF-overexpressing myotubes compared to the controls (Fig. 2G-I). However, large MAFs did not increase—and, in some cases, reduced—the transcription levels of *MYH7*, *MYH2*, and *MYH1*. These findings demonstrate that the overexpression of large MAFs can strongly induce *MYH4* expression in human skeletal muscle cells.

We isolated primary bovine MuSCs and conducted adenoviral infection in bovine myotubes to further confirm that the mechanism underlying *MYH4* expression by large MAFs is conserved across large mammalian species (Fig. S1A). Bovine MuSCs formed multinucleated myotubes with well-defined sarcomere structures after 6 days of differentiation (Fig. S1B). After adenoviral infection, we performed qPCR analysis to evaluate *MYH4* and *MYH7* expression in bovine myotubes (Fig. S1C-H). *MYH4* was barely detectable in bovine myotubes expressing mCherry alone. However, its transcripts were substantially upregulated in all large MAF-overexpressing cells (Fig. S1C-E). Additionally, *MYH7* expression was significantly decreased in all large MAF-overexpressing bovine myotubes (Fig. S1F-H). Given the highly conserved MARE sites in the *MYH4* promoter across species, these findings demonstrate that large MAFs act as universal regulators of *MYH4* expression across mammalian species.

Previous reports suggest that human extraocular muscles exhibit detectable levels of *MYH4* mRNA but not the corresponding protein (*22*, *30*). Considering this notion, two possibilities arise: insufficient *MYH4* mRNA levels or translational suppression of *MYH4* in human skeletal muscle. Thus, we investigated whether *MYH4* mRNA induced by large MAFs can be translated into protein. RNA bound to active ribosomes was extracted using the AHARIBO RNA system (*31*), and RT-qPCR was subsequently performed to assess the binding of *MYH4* mRNA to active ribosomes (Fig. 3A). We detected amplification of *MYH4* mRNA bound to active ribosomes in large MAF-overexpressing myotubes (Fig. 3B, C), whereas no *MYH4* amplification was observed in control myotubes (Fig. 3B, C). To further confirm the presence of the MYH4 protein, we conducted LC-MS/MS analysis in human myotubes overexpressing large MAFs (Fig. 3D). Human MAFA, MAFB, and MAF peptides were only observed in myotubes overexpressing *MAFA*, *MAFB*, and *MAF*, respectively (Fig. 3E-G). Furthermore, *MYH4*-specific peptides were detected in myotubes overexpressing *MAFB* and *MAF* but not in the control (Fig. 3H, I). MYH4 peptides were observed in only one sample of *MAFA*-overexpressing myotubes (Fig. 3H, I), which was consistent with mild *MAFA*-mediated induction of *MYH4* mRNA compared to *MAFB* and *MAF* (Fig. 2G). These findings provide the first evidence of MYH4 in human skeletal muscle cells.

**Fig. 3.**
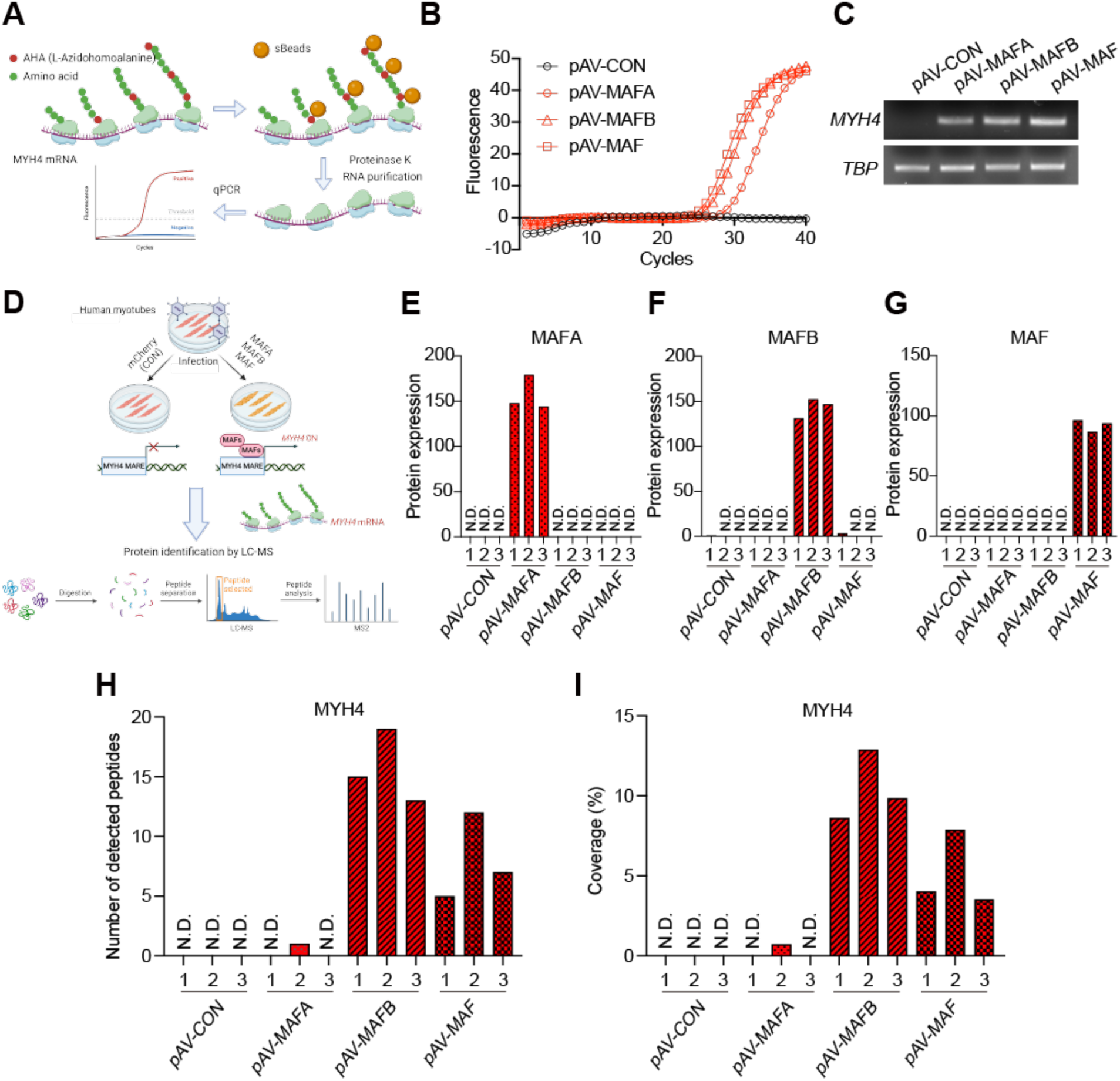
MYH4 mRNA induced by large MAFs is translated into protein in human myotubes. (**A**) A schematic diagram illustrating the protocol for isolating active ribosomes to detect MYH4 mRNA translation in human myotubes following adenoviral overexpression of MAFA, MAFB, or MAF. Created in BioRender. Ryo, F. (2025) https://BioRender.com/g05y857. (**B**) Amplification curve of MYH4 mRNA bound to active ribosomes in human myotubes, as determined using RT-qPCR. (**C**) Gel electrophoresis of qPCR products corresponding to MYH4 and TBP mRNAs bound to active ribosomes, as analyzed in panel (**B**). (**D**) A schematic diagram illustrating the protocol for the detection of MYH4 using LC-MS/MS in human myotubes following adenoviral overexpression of MAFA, MAFB, or MAF. Created in BioRender. Ryo, F. (2025) https://BioRender.com/f96f475. (**E-G**) Protein expression of MAFA (E), MAFB (F), MAF (**G**) in human myotubes following adenoviral overexpression of MAFA, MAFB, or MAF, by LC-MS/MS. mCherry-only was used as the control (pAV-CON) (n = 3/group). N.D.; not detected. (**F**) Number of MYH4-specific peptides detected in human myotubes following adenoviral overexpression of MAFA, MAFB, or MAF, as determined using LC-MS/MS. mCherry-only was used as the control (pAV-CON, n = 3/group). N.D.; not detected. (**G**) Percent peptide coverage of MYH4 in LC-MS/MS analysis. Coverage is expressed as the percentage of the amino acid sequence detected via LC-MS/MS analysis.

### Large MAFs as Key Regulators of Type IIb Myofiber-Specific and Glycolytic Gene Programs in Human Skeletal Muscle Cells

In the present study, we extend our previous findings in mice (*26*) to humans and bovines, providing evidence that *MYH4* is directly controlled by large MAFs in these species. While MYH4 serves as the most critical marker of type IIb myofibers, its expression alone is insufficient to define the formation of authentic type IIb myofibers. To investigate the extent to which large MAFs regulate genes beyond *MYH4*, including genes highly enriched in type IIb myofibers and those involved in glycolysis, we conducted a comprehensive RNA-seq analysis in human myotubes overexpressing *MAFA*, *MAFB*, or *MAF* using the adenoviral vectors described in Fig. 2A.

We observed robust transcriptional changes in myotubes overexpressing each large MAF, with 3,685 transcripts altered in *MAFA*-overexpressing myotubes (1,907 upregulated and 1,778 downregulated), 2,289 transcripts in *MAFB*-overexpressing myotubes (1,195 upregulated and 1,094 downregulated), and 4,446 transcripts in *MAF*-overexpressing myotubes (2,262 upregulated and 2,184 downregulated) compared to control myotubes (Fig. 4A).

**Fig. 4.**
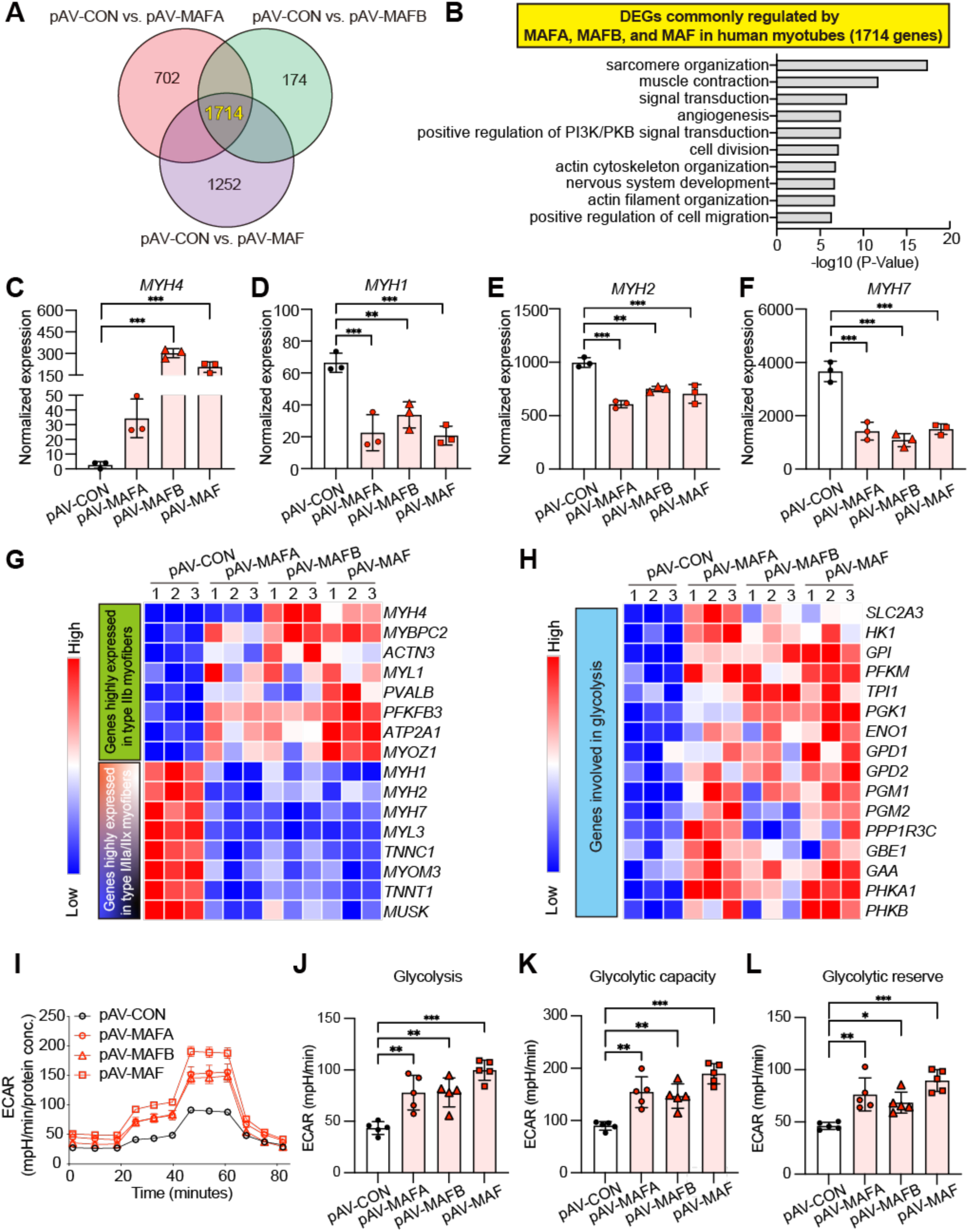
Large MAFs specifically upregulate the expression of type IIb myofiber-associated and glycolytic genes, consequently enhancing glycolysis in human myotubes. (**A**) Venn diagrams showing the differentially expressed genes (DEGs) identified in comparisons between pAV-CON and pAV-MAFA (3,685 genes), pAV-CON and pAV-MAFB (2,289 genes), and pAV-CON and pAV-MAF (4,446 genes). In total, 1,714 DEGs were commonly altered by the overexpression of large MAFs. (**B**) Gene Ontology (GO) analysis of the 1,714 commonly altered genes identified in panel (**A**). (**C-F**) RNA-seq analysis of MYH4, MYH1, MYH2, and MYH7 expression in human myotubes transduced with pAV-CON, pAV-MAFA, pAV-MAFB, or pAV-MAF (n = 3/group). (**G**) A heatmap visualizing the expression of type IIb myofiber-associated genes and other myofiber type-associated genes (n = 3/group). (**H**) A heatmap visualizing the expression of genes involved in glycolysis (n = 3/group). (**I**) Extracellular acidification rate (ECAR) in human myotubes transduced with pAV-CON, pAV-MAFA, pAV-MAFB, or pAV-MAF, as measured using the Seahorse XFe24 Analyzer (n = 5/group). (**J-L**) Glycolysis, glycolytic capacity, and glycolytic reserve calculated from the ECAR data shown in panel (**I**). P values were calculated using Tukey’s test (**C-F**, **J-L**); *P < 0.05, **P < 0.01, ***P < 0.001, **** P < 0.0001.

The expression of 1,714 genes was commonly regulated across myotubes overexpressing large MAFs, including 845 and 869 genes with upregulated and downregulated expression, respectively (Fig. 4A). Gene ontology (GO) analysis of these differentially expressed genes revealed strong enrichment of pathways related to sarcomere organization and muscle contraction (Fig. 4B). Moreover, GO analysis of genes specifically altered by each large MAF in human myotubes suggested unique roles for each transcription factor. However, this may be unrelated to myofiber determination, as no significant enrichment for muscle-related GO terms was detected in these gene sets (Fig. S2A-C). The changes in the expression of adult *Myh* genes closely mirrored the RT-qPCR results shown in Fig. 2G–I. This confirms that *MYH4* expression was specifically and substantially upregulated by large MAFs (Fig. 4C). Its induction was more pronounced in *MAFB*-and *MAF*-overexpressing myotubes than in *MAFA.* Conversely, large MAFs suppressed *MYH7, MYH1,* and *MYH2* expression (Fig. 4D–F).

Consistent with the changes induced by large MAFs in sarcomere organization and muscle contraction (Fig. 4B), the expression of isoforms of contractile proteins enriched in type IIb muscles was markedly upregulated (e.g., *MYBPC2*, *ACTN3, MYL1*) (Fig. 4G). In contrast, the expression of isoforms associated with type I, IIa, and IIx muscles (e.g., *TNNC1*, *TNNT1*) was downregulated (Fig. 4G). Analysis of genes associated with myofiber type transitions revealed that the myocyte enhancer factor 2 (MEF2) family (*MEF2A*, *MEF2B*, and *MEF2C*), which promotes slow oxidative myofibers, was downregulated in large MAF-overexpressing myotubes compared to mCherry-overexpressing controls (Fig. S3A). Transcription factors associated with fast glycolytic muscle genes, including *SIX1*, *SIX4*, *TBX15*, *FNIP1*, *EYA1*, and *SMARCD3* (*8*, *32–36*), were not notably induced by large MAFs in human myotubes (Fig. S3A). Although myotube width in human myotubes did not differ significantly among groups (Fig. 2D), the expression of genes involved in myogenesis, such as *MYOG* (*37*), was downregulated by large MAF overexpression (Fig. S3B).

Type IIb myofibers are further characterized by their high glycolytic gene expression. The loss of large Mafs affects not only fiber type composition but also metabolic profiles in skeletal muscles in mice (*26*, *38*) We sought to determine whether the overexpression of large MAFs elevated the expression of glycolytic genes (*39*) in human myotubes. RNA-seq analysis revealed significant upregulation of glycolytic gene expression in large MAF-overexpressing myotubes (Fig. 4H). Moreover, we confirmed the presence of MARE sites within the promoters of the glycolytic genes listed in Figure 4H.

The profound transcriptional changes in glycolytic genes prompted further investigation into glycolytic activity in human myotubes upon large MAF overexpression. We measured the extracellular acidification rate (ECAR) using the Agilent XF Seahorse technology to generate live-cell bioenergetic profiles. The overexpression of large MAFs significantly enhanced glycolytic rates, including glycolysis, glycolytic capacity, and glycolytic reserve, in large MAF-overexpressing human myotubes compared to controls (Fig. 4I-L). Our findings demonstrate that large MAFs broadly regulate the fast-glycolytic gene program, consequently driving the acquisition of type IIb myofiber-like functional characteristics in human skeletal muscle.

### Large MAF Expression Levels Determine MYH4 expression in Human Skeletal Muscle Tissue

To confirm the association between large MAF and *MYH4* expression in human skeletal muscle tissue, we analyzed *MAFA*, *MAFB*, and *MAF* expression, as well as the combined expression of total large MAFs, using RNA-seq data and myofiber type proportions obtained from muscle biopsies of 291 Finnish participants (men n = 166, women: n = 125) (Fig. 5A, B) and 24 men Russian participants (Fig. 5C, D). In the Finnish cohort, we observed a clear positive correlation between the expression levels of *MAFA*, *MAF*, and total large MAFs with the percentage of fast type II myofibers regardless of sex (Fig. 5A). However, *MAFB* expression showed no significant correlation with the percentage of type II myofibers (Fig. 5A). Additionally, total large MAF expression levels were positively correlated with fast *MYH4* and *MYH1* (Fig. 5B); however, *MYH4* expression levels were markedly low. In contrast, total large MAF expression levels were negatively correlated with *MYH7* expression (Fig. 5B). Furthermore, analysis of each large MAF association with *MYH* genes revealed that *MAFA* and *MAF* showed a positive correlation with *MYH4* (Fig. S4A) and *MYH1* (Fig. S4B) and a negative correlation with *MYH7* (Fig. S4D). The expression levels of *MAFB* did not correlate with those of *MYH* genes, except for *MYH2* (Fig. S4C).

**Fig. 5.**
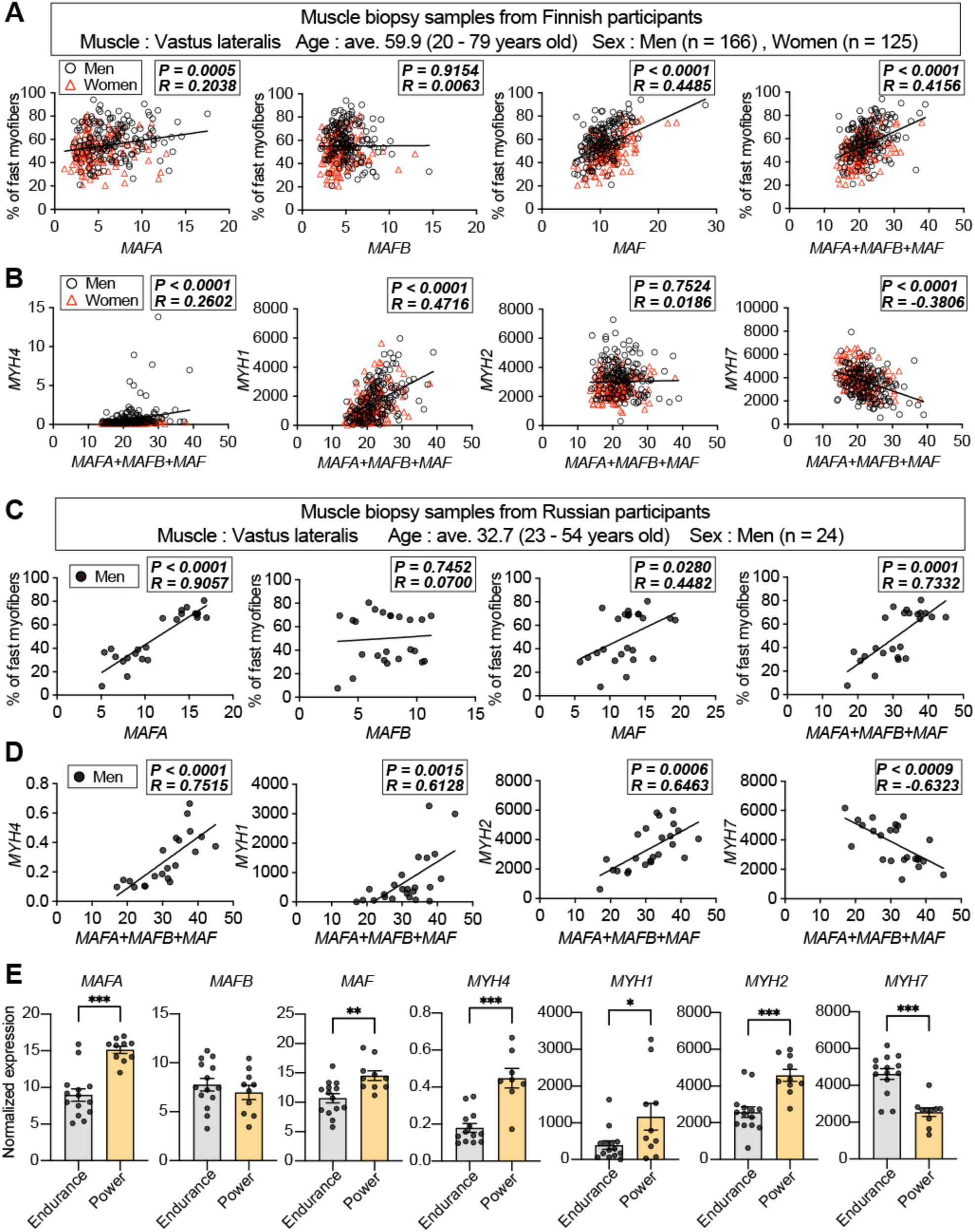
Analysis of human skeletal muscle biopsy samples reveals a positive correlation between MYH4 and large MAF expression, highlighting the potential functionality of large MAFs in human skeletal muscle tissue. (A-B) Muscle biopsy samples were obtained from the vastus lateralis of 291 individuals in Finland, comprising 166 men and 125 women aged 20–79 years (mean age: 59.9 years). (**A**) Scatter plots showing the percentage of fast-twitch myofibers (y-axis) against the mRNA expression levels of *MAFA*, *MAFB*, *MAF*, or total large MAFs (*MAFA*+*MAFB*+*MAF*) (x-axis) for each participant, as determined using RNA-seq analysis. (**B**) Scatter plots showing the mRNA expression levels of *MYH4*, *MYH1*, *MYH2*, or *MYH7* (y-axis) against the total mRNA expression levels of large MAFs (*MAFA*+*MAFB*+*MAF*) (x-axis) for all 291 participants. (**C-D**) Muscle biopsy samples were obtained from the vastus lateralis of 24 men in Russia. The ages of the participants ranged from 20 to 45 years (mean age: 32.7 years). (**C**) Scatter plots showing the percentage of fast-twitch myofibers (y-axis) against the mRNA expression levels of *MAFA*, *MAFB*, *MAF*, or total large MAFs (*MAFA*+*MAFB*+*MAF*) (x-axis) for each participant, as determined through immunohistochemical and RNA-seq analyses. (**D**) Scatter plots showing the mRNA expression levels of *MYH4*, *MYH1*, *MYH2*, or *MYH7* (y-axis) against the total mRNA expression levels of large MAFs (*MAFA*+*MAFB*+*MAF*) (x-axis) for all 24 participants. (**E**) RNA-seq analysis of MAFA, MAFB, MAF, MYH4, MYH1, MYH2, and MYH7 expression in Russian muscle biopsy samples, categorized by training type: endurance training (n=14) and power training (n=10). P values were calculated using a two-sided Pearson correlation test (**A-D**) and Student’s *t*-test (**E**); *P < 0.05, **P < 0.01, ***P < 0.001, R; Pearson correlation coefficient. Outliers were identified using Grubbs’ test (Alpha = 0.05); three participants with outlying *MYH4* expression levels were excluded from the analysis (**D-E**).

Similar correlation patterns were obtained in the muscle biopsy data from Russian samples compared to the Finnish dataset (Fig. 5C, D). Specifically, we detected a clear positive correlation between the expression levels of *MAFA*, *MAF*, and total large MAFs with the percentage of fast type II myofibers in the muscle samples of Russian participants, as quantified through immunostaining (Fig. 5C). Consistent with the samples obtained from Finnish participants, *MAFB* showed no correlation with the percentage of type II myofibers in the Russian dataset (Fig. 5C). Furthermore, total large MAF expression levels showed the strongest positive correlation with fast MYH (*MYH4*, *MYH1*, and *MYH2*) but was negatively correlated with the slow MYH, *MYH7* in the Russian cohort (Fig. 5D). Analysis of individual large MAF associations with *MYH* genes revealed a similar trend: *MAFA* and *MAF* expression levels, but not *MAFB*, were positively correlated with *MYH4*, *MYH1*, and *MYH2* and negatively associated with *MYH7* (Fig. S5A-D). Although *MYH4* expression levels were considerably low across all samples compared to other *MYH* genes, our comprehensive analysis further reinforces the conclusion that large MAFs regulate *MYH4* expression in human skeletal muscle *in vivo*.

Lastly, to investigate the responsiveness of large MAFs to exercise training inducing myofiber type shifts, we examined their expression under different training regimens. A Russian dataset documenting individual physical activity was categorized into two groups: endurance-trained and power-trained participants. RNA-seq analysis revealed that *MAFA* and *MAF* expression were significantly higher in power-trained participants compared to endurance-trained participants, while *MAFB* expression remained comparable between the groups (Fig. 5E). Similarly, the expression of fast *MYH* genes, including *MYH4*, was upregulated in power-trained participants, whereas slow *MYH7* expression was elevated in endurance-trained participants (Fig. 5E). These findings suggest that *MYH4* expression is influenced by exercise type, with power training enhancing *MAFA* and *MAF* expression in human skeletal muscles, and highlight large MAFs as potential biomarkers for assessing muscle functionality in athletes.

## Discussion

The limb skeletal muscles of small animals such as mice and rats predominantly comprise fast-twitch myofibers, including type IIa (*Myh2)*, IIx (*Myh1*), and IIb (*Myh4*). In contrast, human skeletal muscle lacks type IIb myofibers (*MYH4*), the fastest glycolytic myofiber type in the limb muscles of mammals (*2*, *4*). In the present study, building on our previous findings that the large Mafs are a principal regulator of type IIb myofiber specification in mice (*26*), we demonstrates that large MAFs specifically and robustly regulate the expression of type IIb myofiber-specific genes, including *MYH4*, in human and bovine skeletal muscle cells. These results challenge the long-held belief that *MYH4* expression and type IIb myofibers are evolutionarily lost in large mammals. To our knowledge, this study is the first to show that large MAFs drive a substantial upregulation not only *MYH4* mRNA but also MYH4 protein in human skeletal muscle cells. Despite the relatively low expression of *MYH4* compared to *MYH1* and *MYH2*, large MAF levels positively correlated with the proportion of fast myofibers and fast myosin expression, including that of *MYH4*. Our findings suggest that large MAFs may unlock latent potential for the development of type IIb myofibers in humans, potentially enhancing rapid muscle contraction and power generation.

Our previous work revealed that large Mafs (*Mafa, Mafb,* and *Maf*) specifically bind to the MARE sequence in the *Myh4* promoter, which is essential for the formation of type IIb myofibers in mice (*26*). As large mammals, including horses, bovines, and humans, have lost type IIb myofibers and *MYH4* expression for reasons that remain unclear, we hypothesized that a decrease in large MAF expression may be a key contributor to the loss of *MYH4* in these species. The low expression of *MYH4* in large animals was associated with the expression level of large MAFs in the present study, suggesting that the levels of large MAFs partly determine the extent of *MYH4* expression. This conclusion raises another question: Why is large MAF expression reduced in large animals such as humans?

Large MAF activity is regulated through multiple mechanisms, including transcriptional and translational controls, as well as regulation by microRNAs (*40*). *MyoD* is a potential regulator of mouse *Mafb* and *Maf* promoter activity, as shown by luciferase assays (*41*, *42*). Additionally, MYOD binds to the *Mafa* promoter region in C2C12 cells, a mouse myoblast cell line (data not shown). However, *MYOD* expression was not downregulated in human myotubes in the present study (Fig. S3B). This suggests that the low expression of large MAFs in human skeletal muscle cannot be attributed to *MYOD* dynamics alone. A novel mechanism underlying the regulation of *Maf* expression was recently proposed in mice; *Maf* expression was significantly reduced in mice lacking *Cacna1s*, which encodes the α1s subunit of Cav1.1 (*38*). As the rapidity of calcium transients varies among animal species (*43*), the diversity of calcium signaling or differences in *Cacna1s* expression across species may affect *Maf* expression levels.

Post-transcriptional regulation of large MAFs may partially explain the reduced expression of large MAF mRNA and protein in humans. miR-155 and miR-1290 interact with the 3′UTR of *Maf* in laryngeal squamous cell carcinoma (*40*, *44*, *45*). miR-1290 is exclusively found in great apes, including humans (*40*, *46*). This demonstrates the possibility of human-specific mechanisms that suppress *Maf* expression, such as those mediated by miR-1290. Additionally, epigenetic modifications, such as differences in histone acetylation or DNA methylation at the *Maf* locus, may contribute to the lower expression of *Maf* observed in humans and other large mammals. However, further investigations are needed to elucidate these potential mechanisms.

In addition to the low levels of large MAFs in human skeletal muscle, other suppressive mechanisms regulating *MYH4* may exist in humans. Nucleotide differences in the *MYH4* promoter between humans and mice may reduce transcriptional activity by disrupting the binding sites for MEF2 and serum response factor (*47*, *48*). While this hypothesis provides an interesting perspective, our results challenge its relevance. Specifically, all *MAFA*, *MAFB*, and *MAF* markedly upregulated *MYH4* expression in human myotubes, despite the reported differences in the *MYH4* promoter region. Furthermore, the MAF recognition element (MARE), a key binding site for large MAFs, was highly conserved across animal species (Fig. 1B). This result confirms that the MARE region in the human *MYH4* promoter retains its functional relevance regardless of the nucleotide differences highlighted in the previous study. Thus, large MAFs can likely override the effects of such promoter differences, directly activating *MYH4* expression in human myotubes. Therefore, we propose that the nucleotide differences in the *MYH4* promoter are unlikely to fully account for the low expression of *MYH4* in humans.

An fMyh-SE region located 42 kb upstream of the *Myh2-Myh1-Myh4* fast myosin gene cluster was recently identified as a critical regulator of fast myosin gene expression in mice (*25*, *49*). Dos Santos et al. (*25*) and Sakakibara et al. (*49*) demonstrated that the fMyh-SE region interacts exclusively with each *fMyh* promoter through a 3D chromatin looping structure, enabling precise and robust expression of specific fast myosin genes. Although our previous and present studies have identified large MAFs as specific transcriptional regulators of *Myh4* (*MYH4*) in both mice and humans, the transcription factors specifically regulating *Myh1* and *Myh2* remain unknown. Given the exclusive regulation mediated by fMyh-SE, we hypothesize that transcription factors specific to *MYH1* or *MYH2* preferentially dominate the fMyh-SE region in humans. This dominance could restrict access to *MYH4*, thereby contributing to its low expression in human skeletal muscle. However, our findings suggest that overexpression of large MAFs can override this exclusivity. Large MAFs may “hijack” the fMyh-SE region, facilitating *MYH4* expression even in human skeletal muscle, where endogenous large MAF expression is low. This model highlights the complex interplay between super-enhancer regulation and transcription factor specificity during the regulation of myosin gene expression across species, which partially explains why human skeletal muscle has lost *MYH4* expression.

Some previous studies have reported that *MYH4* mRNA can be detected in a small subset of specialized muscles, such as the human masseter and laryngeal muscles, as well as in skeletal muscle from patients with Duchenne muscular dystrophy. However, no detectable MYH4 protein was observed in previous studies (*22*, *23*, *30*, *47*). The present study provides the first evidence of MYH4 protein expression via the overexpression of large MAFs in human skeletal muscle.

Type IIb myofibers, which are characterized by their fast contraction speeds and high force generation, also have the highest concentration of glycolytic enzymes compared to other type II myofibers. As a limitation, the present study did not determine whether the induction of type IIb myofibers is beneficial or detrimental to human skeletal muscle or overall health. However, our reanalysis of snRNA-seq in human skeletal muscle (*16*) demonstrated that fast myofiber signatures, including fast MYH, are preferentially lost with aging, along with a decline in *MAF* expression. Furthermore, shifts in myofiber types, whether in pathological or physiological contexts, can influence the skeletal muscle microenvironment, including vasculature, the extracellular matrix, the stem cell niche, and MuSCs. Thus, we hypothesize that restoring MAF expression could counteract the pathological features of sarcopenia by improving muscle mass and function, and by rejuvenating the microenvironment of aged skeletal muscle. This could include enhancing MuSC-mediated regeneration, ultimately providing a comprehensive approach for mitigating the detrimental effects of aging on skeletal muscle.

Large MAFs specifically activated type IIb-specific genes and glycolytic-associated gene programs, suggesting that they exhibit some functional redundancy in inducing these gene sets. They commonly downregulated the expression of members of the MEF2 family (*MEF2A, MEF2B,* and *MEF2C*) that contribute to the transcription of slow oxidative myofiber-specific gene programs (*50–53*). Furthermore, the overexpression of *Smarcd3*, which encodes BAF60c, in mouse skeletal muscle strongly promotes glycolytic gene expression and enhances insulin sensitivity (*36*). In contrast, large MAFs downregulated *SMARCD3* expression in human myotubes in our study. This suggests that large MAFs regulate glycolytic genes independently of *SMARCD3*. Consistent with this, we identified MARE sequences in the promoters of all glycolysis-related genes listed in Fig. 4H, indicating the direct regulatory role of large MAFs in glycolytic gene activation.

Our RNA-seq analysis of myotubes overexpressing each large MAF also revealed distinct functional signatures for each transcription factor. For instance, pathways related to the ubiquitin-dependent protein catabolic process and protein transport were specifically enriched in MAFA-overexpressing myotubes, whereas pathways associated with interferon beta production and innate immune response were enriched in MAFB-overexpressing myotubes (Fig. S2A, B). Cytoplasmic translation and ribosomal biogenesis pathways were prominently enriched in MAF-overexpressing myotubes (Fig. S2C). Although MAFA induced *MYH4* expression to a lesser extent than MAFB or MAF (Fig. 2G-H, Fig. 4C), it equally promoted the expression of most glycolytic genes (Fig. 4H). These findings indicate that large MAFs have unique targets and preferences despite their shared role in the activation of type IIb-specific and glycolytic gene programs. Therefore, future studies should explore co-transcriptional factors and chromatin remodeling complexes that interact with each large MAF to further explore how large MAFs orchestrate diverse gene programs in mice and human skeletal muscle.

Our *in vitro* findings were further validated *in vivo* using RNA-seq data from human muscle biopsy samples. Despite the low expression levels of *MYH4*, a clear positive correlation was observed between *MYH4* expression and the expression of *MAFA*, *MAF*, or total Large MAFs across two independent cohorts. The expression of *MYH1*, which encodes type IIx myosin, was also positively correlated with *MAFA, MAF,* and total large MAFs in both cohorts. Unlike mouse and rat skeletal muscle, human type IIx myofibers exhibit the fastest contraction speeds and the highest glycolytic activity, while showing the lowest succinate dehydrogenase activity (*2*, *17*, *54*). In the present study, large MAFs did not directly induce *MYH1* expression but strongly upregulated the expression of glycolytic genes. Therefore, we hypothesize that the observed correlation between *MYH1* and large MAF expression reflects the role of large MAFs in driving glycolytic gene programs, specifically in fast-glycolytic fibers, such as type IIx myofibers. Thus, large MAFs may indirectly associate with *MYH1* expression by activating glycolytic gene programs in human skeletal muscles.

Resistance training, rather than endurance training, was associated with increased expression of *MAFA* and *MAF* in the Russian dataset, alongside the upregulation of fast *MYH* genes, including *MYH4*. This finding raises the possibility that specific resistance training regimens could modulate MAFA and MAF expression to enhance MYH4 levels in human athletes. Interestingly, ACTN3, a key gene associated with athletic speed, showed the strongest positive correlation with MAF expression in humans. Given that the ACTN3 promoter contains a MARE site, MAFA and MAF expression levels may serve as potential biomarkers for guiding individuals toward sports that align with their muscle fiber characteristics. Furthermore, our findings suggest that certain individuals, such as professional athletes, may naturally exhibit higher levels of large MAFs or possess gain-of-function genetic variants that promote the development of type IIb-like muscle traits. Future studies should investigate these genetic variants across diverse populations to elucidate their role in athletic performance and muscle fiber specialization.

In conclusion, our previous and current studies have firmly established large MAFs as key regulators of fast glycolytic myofiber programs across mammalian species, including humans. Furthermore, our findings highlight large MAFs as novel biomarkers for assessing fast myofiber proportions, and as predictive markers for athletic adaptation or training responsiveness. This demonstrates the potential applicability of large MAFs in sports science. Importantly, fast-twitch myofibers are preferentially lost during aging, potentially due to a decline in the expression of large MAFs. Our work provides the exciting possibility of reintroducing fast-glycolytic type IIb myofibers—long absent in humans—through the manipulation of large MAFs or their upstream pathways. This approach may provide a novel strategy for restoring fast-twitch myofibers in aged skeletal muscle.

## Materials and Methods

### Human iPS Cloning and Generation of Muscle Stem Cells

The healthy donor-derived human pluripotent stem cell (hiPSC) clone 414C2 (*55*) was used to harvest iPSC-derived muscle stem cells (iMuSCs) after myogenic differentiation. Myogenic induction of hiPSCs to obtain iMuSCs was performed as described in our previous study (*56*). Briefly, undifferentiated hiPSCs were plated onto a 6-well plate coated with perlecan-binding laminin421E8 fragment (p421E8) in StemFit AK02N (Ajinomoto) and 10 μM Y-27632 (Nacalai) at 5 × 10^3^ cells/well. After three days, the medium was replaced with CDMi supplemented with 10 μM CHIR99021 (CHIR, Wako) and 5 μM SB431542 (SB, Wako). CDMi is composed of IMDM (Wako) and Ham’s F-12 medium (Wako) (ratio, 1:1) supplemented with 1% bovine serum albumin (BSA) (Sigma-Aldrich, Burlington, MA, USA), 1% penicillin-streptomycin mixed solution (Nacalai), 1% CD lipid concentrate (Thermo Fisher Scientific, Waltham, MA, USA), 1% insulin-transferrin selenium (Thermo Fisher Scientific), and 450 μM 1-thioglycerol (Sigma-Aldrich).

After seven days of differentiation, the cells were dissociated with Accutase (Nacalai) and plated onto a p421E8-coated 6-well plate (4.5 × 10^5^ cells/well) in CDMi medium supplemented with 10 μM CHIR, 5 μM SB, and 10 μM Y-27632. On differentiation day 14, the cells were dissociated with Accutase and placed onto an iMatrix-511 (Nippi)-coated 6-well plate (8 × 10^5^ cells/well) in CDMi medium supplemented with 10 μM Y-27632. The medium was replaced with serum-free culture medium (SF-O3; Sanko Junyaku) supplemented with 0.2% BSA, 200 μM 2-2-mercaptoethanol (2-ME), 10 ng/mL IGF-1 (PeproTech), 10 ng/mL recombinant human bFGF (Oriental Yeast), and 10 ng/mL recombinant human HGF (PeproTech) after three days. The medium was changed twice a week until day 38 of differentiation. On differentiation day 38, it was replaced with 2% horse serum (HS, Sigma-Aldrich) in DMEM (Nacalai) supplemented with 200 μM 2-ME (Nacalai), 10 ng/mL IGF-1, 5 μM SB, 0.5% penicillin-streptomycin mixed solution (Nacalai), and 2 mM L-glutamine (Nacalai). Thereafter, the medium was changed every 2– 3 days, and the cells were cultured in the medium until weeks 11–12 of differentiation.

### Isolation of iMuSCs through Flow Cytometric Sorting

The cells were harvested for flow cytometric sorting after 11–12 weeks of differentiation (*57*). Briefly, cells were dissociated with Collagenase mix solution at 37 °C for 7 min. The collagenase mix solution is composed of DMEM supplemented with 0.05% Collagenase H (Meiji Seika Pharma) and 0.5% Collagenase G (Meiji Seika Pharma). Accutase was added to the solution and incubated at 37 °C for 10 min. The solution was subsequently neutralized with DMEM supplemented with 10% HS and centrifuged at 380 × *g* for 7 min at 4 °C. Next, the supernatant was removed. Cells were then resuspended in DMEM supplemented with 10% HS and filtered through a 40-µm cell strainer (Corning). Density gradient centrifugation was performed using Optiprep (Serumwerk Bernburg) in DMEM supplemented with 10% HS (ρ = 1.11 g/mL) according to the manufacturer’s instructions to remove debris and dead cells. The fraction of living cells was resuspended in HBSS (Thermo Fisher Scientific) and 1% BSA before incubation with an APC-conjugated anti-CDH13 antibody (5 × 10^6^ cells/mL) on ice for 15 min (*28*). Flow cytometry was performed using Aria II (BD) in accordance with the manufacturer’s instructions. Gating of the CDH-13-positive fraction was determined using unstained cells as the baseline control. After sorting the CDH-13-positive cells, the suspension was centrifuged at 780 × *g* for 10 min at 4 °C. The supernatant was removed, the 2 × 10^5^ sorted cells were cryopreserved in 200 μL Bambanker (NIPPON Genetics).

### Re-culture and Myogenic Differentiation of Sorted iMuSCs

Frozen sorted cells were rapidly thawed in a water bath at 37 °C. They were then suspended in a growth medium containing 39% DMEM (Gibco), 39% Ham’s F12, 20% FBS (Gibco), and 1% UltroserG (Pall Life Sciences). Next, the cells were centrifuged at 400 × *g* for 10 min at 4 °C, as described previously (*58*, *59*). After removal of the supernatant, the cells were resuspended in a growth medium and plated on to an iMatrix-coated (nippi) 12-well plate (2 × 10^4^ cells/well). The medium was replaced with a differentiation medium containing 2% HS in DMEM after three days. Thereafter, the medium was changed every 2 days until analysis. The growth and differentiation media were supplemented with 1% penicillin-streptomycin.

### Preparation and Culture of Primary Bovine Muscle Stem Cells

The thoracic longissimus muscle was excised from a 22-month-old Holstein cow. Primary bovine muscle stem cells were isolated as described previously, with some minor modifications (*60*, *61*). Connective tissue, blood vessels, and fat tissue were removed from isolated muscle tissue under dissection microscopy. Minced muscle pieces were treated with 0.2% collagenase type II (Worthington Biochemical Corporation) in HBSS (Thermo Fisher) for 30–40 min at 37 °C. Muscle slurry was homogenized with an 18G needle passing through 10 times and subsequently incubated at 37 °C for another 10 min. Debris was filtrated through 100-µm and 40-µm cell strainers (Corning Japan KK). The obtained cells were cultured on non-coated dishes in 20% FBS (Biowest) in F10 (Thermo Fisher) supplemented with 2.5 ng/mL bFGF (Nacalai Tesque, Inc.) for 60 min. Next, a non-adherent fraction was collected as primary bovine MuSCs and cultured on collagen-coated dishes (AGC Techno Glass Co., Ltd.) for expansion. Cells with a few passages were cryopreserved and used for the experiments. Enrichment of primary bovine MuSCs and myotubes were confirmed through immunostaining with an antibody against sarcomeric-α-actinin, which is specifically expressed in skeletal muscles. The cells generated multinucleated myotubes after myogenic differentiation. In addition, >50% of the cultured cells were positive for sarcomeric-α-actinin, which is specifically expressed in skeletal muscle cells.

Frozen bovine cells were rapidly thawed in a water bath at 37 °C and followed the same protocol as that for iMuSCs. Briefly, bovine cells were resuspended in growth medium and plated on to an iMatrix-coated (nippi) 12-well plate (2 × 10^4^ cells/well). The medium was replaced with a differentiation medium containing 2% HS in DMEM after two days. Thereafter, the medium was changed every 2 days until analysis. The growth and differentiation media were supplemented with 1% penicillin-streptomycin.

### Overexpression of Adenovirus-mediated large MAFs

The following adenoviral vectors were used to overexpress the large MAF transcription factor family in human iMuSCs and primary bovine MuSCs: pAV[Exp]-mCherry-CMV>hMAFA, pAV[Exp]-mCherry-CMV>hMAFB, and pAV[Exp]-mCherry-CMV>hMAF. pAV[Exp]-mCherry-CMV was used for control. All vectors were constructed and packaged by VectorBuilder Inc. (Chicago, IL, USA), with vector IDs as follows: VB010000-9300kfr (mCherry), VB900139-7528hhe (hMAFA), VB900139-7526uvc (hMAFB), and VB900139-7527axe (hMAF). The myotubes were incubated with adenovirus at a multiplicity of infection of 250 in differentiation medium for 48 h two days post-differentiation of human iMuSCs or primary bovine MuSCs. The medium was then replaced with fresh differentiation medium without adenovirus. The myotubes were used for subsequent experiments four days post-infection.

### Immunofluorescence

Cultured cells were washed twice with PBS after removing the media and fixed with 4% paraformaldehyde (PFA) for 15 min. The cells were permeabilized and blocked using 5% goat serum in 0.1% Triton PBS for 1 h at room temperature and then washed 3 times with PBS. The cultured cells were incubated with primary antibodies at 4°C overnight. All immunostained samples were visualized using appropriate Alexa Fluor conjugated secondary antibodies (Thermo Fisher Scientific) for 1 h. Primary antibodies were against RFP (600-401-379, Rockland), α-actinin (A7811; Sigma-Aldrich). Secondary antibodies conjugated with Alexa Fluor-488, Alexa Fluor-555, and Alexa Fluor-647 were used.

### Analysis of Actively Translated mRNA in Human Myotubes using AHA-mediated Ribosomal Isolation

The active ribosome was isolated with the AHARIBO RNA system (Immagina) as described previously (*31*) to analyze MYH4 mRNA translation in human myotubes. Briefly, adenovirus-infected human myotubes were washed once with PBS and treated for 40 min with methionine-free growth medium (Thermo Fisher Scientific) supplemented with 2% FBS and 0.8 mM l-leucine to deplete methionine reserves. After 40 min, 10 µl of AHA reagent was added to the medium. The cells were then incubated at 37 °C for 5 min, following the addition of 2.6 µl sBlock for 5 min at 37 °C. Next, they were placed on ice and washed once with 1 ml of cold PBS. PBS was carefully removed with a pipette, and cells were lysed with 45 µl of cold lysis buffer using a cell scraper. Cell lysate was transferred to a 1.5-ml microcentrifuge tube, and cell debris was pelleted through centrifugation at 20,000 × *g* for 5 min at 4 °C. The supernatant was transferred to a new tube and kept on ice for 20 min. Absorbance was measured by NanoDrop at 260 nm, with lysis buffer as blank subtraction.

Two absorbance units (AU) was transferred to a new tube, and the volume was adjusted to 100 µl with freshly prepared SWB buffer to capture active ribosomal complexes. To this, 100 μl of sBeads was added, and the mixture was incubated for 60 min at 4 °C on a rotating wheel. Next, the supernatant was removed, and the beads were washed twice with WSS buffer. The beads were resuspended in 200 μl of SWB buffer. RNA was then extracted from the resuspended SWB buffer containing the beads. The extracted RNA concentration was measured at 260 nm using a NanoDrop spectrophotometer, and cDNA was subsequently synthesized.

### Quantitative Analysis of Transcripts using Reverse Transcription PCR

Total RNA was extracted from human and bovine myotubes using TRIzol reagent (Life Technologies) according to the manufacturer’s instructions. RNA (500 ng) or 340 ng of active ribosome-associated RNA was reverse transcribed using either the ReverTra Ace Kit with genomic DNA remover (Toyobo, Osaka, Japan) or the QuantiTect Reverse Transcription Kit (Qiagen) for cDNA synthesis. Quantitative PCR was performed using the THUNDERBIRD SYBR qPCR system (Toyobo, Osaka, Japan) on a TP850 Thermal Cycler Dice Real-Time System (Takara Bio, Kusatsu, Japan). Primer sequences are provided in Supplementary Table 1. Transcription levels were normalized to the TATA-box-binding protein (TBP) transcript levels for each sample.

### RNAscope

Adenovirus-infected human myotubes were fixed in 4% paraformaldehyde for 15 min to visualize MYH4 transcripts. Fixed cells were processed using the RNAscope Multiplex Fluorescent V2 Assay (Advanced Cell Diagnostics, 323270), with MYH4 transcripts hybridized to Hs-MYH4-C1 probes (ADC, 1583381-C1). Briefly, fixed cells were washed twice with PBS before incubation for 10 min in hydrogen peroxide solution. Cells were then treated with protease III and diluted 1:15 in PBS for 10 min. Next, they were washed twice with and hybridized with the Hs-MYH4-C1 probe for 2 h at 40 °C. MYH4 transcripts were visualized using the TSA Vivid Fluorophore Kit 520 (R&D Systems) according to the manufacturer’s instructions. Cells were washed twice in PBS, and immunofluorescence analysis was performed using antibodies against α-actinin and RFP, as described above. Nuclei were counterstained with DAPI.

### RNA-seq Analysis of Human iMuSC-derived Myotubes

Total RNA was extracted from human myotubes on day 4 of infection using TRIzol reagent (Thermo Fisher Scientific). The RNA-seq library was prepared using the NEBNext Ultra Directional RNA Library Prep Kit (New England Biolabs, Ipswich, MA, USA) after ribosomal RNA (rRNA) depletion (NEBNext rRNA Depletion Kit; New England Biolabs). Paired-end (2 × 36 bases) sequencing was performed using the NextSeq500 platform (Illumina, San Diego, CA, USA). FASTQ files were imported to the CLC Genomics Workbench (Version 10.1.1; Qiagen, Hilden, Netherlands). Sequence reads were then mapped to the human reference genome (hg19). Gene expression was calculated as total read counts normalized by transcripts per million. Genes with zero counts in any sample were excluded, and differential expression was analyzed using the Empirical Analysis of DGE tool (edgeR test) in the CLC Main Workbench (Version 22.0; Qiagen). DEGs were extracted among conditions (pAV-mCherry vs. pAV-MAFA vs. pAV-MAFB vs. pAV-MAF) with a false discovery rate–corrected P < 0.05. The DAVID Bioinformatics Resources 6.8 (*62*) was used for GO analysis, with P-value < 0.05.

### Seahorse XF Glycolysis Stress Test

iMuSCs were seeded in XFe 24-well plates (Seahorse Bioscience, North Billerica, MA, USA) at a density of 1 × 10⁴ cells/well. Adenoviral infection was performed on iMuSCs after 2 days of differentiation in DM. The ECAR was measured using a Seahorse XFe24 analyzer (Seahorse Bioscience) four days after infection. For the glycolysis stress test, glucose, oligomycin, and 2-deoxyglucose (2-DG) were sequentially added to the wells at final concentrations of 10 mM, 1.0 µM, and 50 mM, respectively. Glucose serves as a substrate for glycolysis, whereas oligomycin inhibits mitochondrial ATP synthase, consequently increasing reliance on glycolysis for energy production. Finally, 2-DG, a competitive inhibitor of glucose, effectively shuts down glycolysis.

### Single-Nucleus RNA-seq Analysis

Single-nucleus RNA sequencing data were downloaded from the Human Muscle Ageing Cell Atlas database (https://db.cngb.org/cdcp/hlma/), and the Seurat pipeline was applied to this dataset. The Seurat objects underwent normalization, scaling, and dimensional reduction. Types I and II myonuclei were classified based on previous studies (*16*). We used density plots generated through the Nebulosa package in our snRNA-seq analysis to assess the localization of large MAFs within myonuclei (*63*). Expression levels of large MAFs and MYH genes were measured in clusters of type II-positive myonuclei. The gene expression in the snRNA-seq data was illustrated using two approaches: first, by plotting the expression levels of the gene across all myonuclei from young (15–46 years old, n = 8) and old (74–99 years old, n = 19) individuals; and second, by comparing the gene expression levels within myonuclei aggregated for each individual, grouped into young and old categories.

### Mass Spectrometry

Human myotube samples (10 mg) were reduced, alkylated, and digested with trypsin/Lys-C using the iST kit (iST 8x, P.O.00001, PreOmics, Germany) according to the manufacturer’s instructions. The samples were then evaporated to dryness *in vacuo* to obtain residues. These residues were dissolved in 10 mL of LC-LOAD buffer from the iST kit. Peptides (200 ng) were injected into nanoLC/ESLI-MS/MS systems [LC: ACQUITY UPLC M-Class system (Waters, MA, USA), MS: ZenoTOF7600 (Sciex)]. The peptide samples were subsequently separated using the nanoEase M/Z Peptide CSH C18 column (1.7 mm, 0.075 3 150 mm) (Waters) at a flow rate of 300 nL/min. Mobile phase A consisted of a 0.1% (v/v) solution of formic acid in water, whereas mobile phase B consisted of a 0.1% (v/v) solution of formic acid in acetonitrile. The linear gradient conditions were as follows: 0–17 min: 2% solvent B, 17–125 min: 30% B, 126–127 min: 40% B, 127–132 min: 80% B, 132–133 min: 90% B, 133–153 min: 90% B, 153–154 min: 2% B, 154–180 min: 2% B. The mass spectrometer was operated in a Zeno SWATH acquisition scheme with 100 variable-size windows, and 25 ms accumulation time was used. The Zeno SWATH raw MS data were processed using DIA-NN 1.8.1,(*64*) available on github (DIA-NN github repository).

The human spectral libraries were generated from the human spectral library and human UniProt database (id UP000005640, reviewed, canonical). The DIA-NN search parameters were as follows: experimental data search enzyme, trypsin; missed cleavage sites, 1; peptide length range, 7–30; precursor mass charge range, 1–4; precursor m/z range, 300–1800; fragment ion m/z range, 200–1800; and static modification, cysteine carbamidomethylation. The protein identification threshold was set at <1% for both peptide and protein FDR. Each peptide assigned to MYH4 was verified through a BLAST search, and peptides specific for MYH4 were counted. The newly acquired mass spectrometry proteomic data have been deposited to the Proteome Xchange Consortium via jPOSTrepo (*65*) with the dataset identifier JPST003487.

### Comparison of Gene Expression Patterns across Multiple Animal Species

To compare MYH4 and large MAF expression scores across animal species, Bgee v15.2.0 was used (*27*). Expression scores were based on the rank of a gene in a given condition according to its expression levels, normalized using the minimum and maximum rank within the species. Expression scores range from 0 to 100, with lower scores indicating lower expression of the gene in the condition compared to other genes. These scores are normalized, making them comparable across genes, conditions, and species. The anatomical entities included in the analysis were skeletal muscle tissue (frog, rat, opossum, dog, and cow), hindlimb and tentacle muscle (mouse and human), and general muscle tissue (horse).

### Motif Enrichment Analysis

The presence of MARE sites in the promoter regions of genes were predicted through the Find Individual Motif Occurrences (FIMO) algorithm of the MEME Suite (*66*), using the position weight matrix available from JASPAR (https://jaspar.genereg.net/). This analysis was run from 3 kb upstream to 3 kb downstream of the *Myh4* transcriptional start in mice, rats, opossum, frogs, humans, dogs, cows, and horses.

### Study Participants

The Russian part of the study was approved by the Ethics Committee of the Federal Research and Clinical Center of Physical–Chemical Medicine of the Federal Medical and Biological Agency of Russia (protocol no. 2017/04). The Finnish part of the study was approved by the Hospital District of Helsinki and Uusimaa (this data was used with permission; Database of Genotypes and Phenotypes (dbGaP) Study Accession: phs001048.v2.p1). Written informed consent was obtained from each participant. The study complied with the Declaration of Helsinki and ethical standards for sport and exercise science research.

The Russian gene expression study (analysis of MAF gene expression in m. vastus lateralis) involved 24 men (mean age ± SD: 32.7 ± 8.9 years; mean height: 180.8 ± 6.8 cm; mean body mass: 80.1 ± 11.6 kg; mean percentage of fast-twitch muscle fibers: 50.3 ± 21.3%; mean percentage of slow-twitch muscle fibers: 52.7 ± 21.6%; cross-sectional area (CSA) of fast-twitch muscle fibers: 5934 ± 1833 μm^2^; CSA of slow-twitch muscle fibers: 5545 ± 1197 μm^2^). Outliers were identified using Grubbs’ test (Alpha = 0.05) in GraphPad Prism; three participants with outlying MYH4 expression levels were excluded from the analysis.

The Finnish study involved 291 individuals (166 men, mean age 59.5 ± 8.1 years; mean height: 176.7 ± 6.7 cm; mean body mass: 87.3 ± 15.1 kg; mean percentage of fast-twitch muscle fibers: 58.9 ± 14.7%; mean percentage of slow-twitch muscle fibers: 41.1 ± 14.7%; 125 women, mean age: 60.3 ± 8.1 years; mean height: 162.8 ± 5.6 cm; mean body mass: 71.8 ± 9.8 kg; mean percentage of fast-twitch muscle fibers: 49.9 ± 13.3%; mean percentage of slow-twitch muscle fibers: 50.1 ± 13.3%) from the FUSION study as previously described (*67*).

### Evaluation of Muscle Fiber Composition in Human Muscle Biopsy Samples

Vastus lateralis samples of Russian participants were obtained from the left leg using the modified Bergström needle procedure, with aspiration under local anesthesia using a 2% lidocaine solution. Samples were frozen in liquid nitrogen and stored at −80 °C before analysis. Serial cross-sections (7 μm) were obtained from frozen samples using an ultratom (Leica Microsystems, Wetzlar, Germany). Sections were then thaw-mounted on Polysine glass slides, maintained at room temperature (RT) for 15 min and incubated in PBS (3 × 5 min). The sections were then incubated at RT in primary antibodies against slow or fast isoforms of the myosin heavy chains (M8421, 1:5000; M4276; 1:600, respectively; Sigma-Aldrich) for 1 h and incubated in PBS (3 × 5 min). Next, the sections were incubated at RT in secondary antibodies conjugated with FITC (F0257; 1:100; Sigma-Aldrich) for 1 h. Antibodies were removed, and the sections washed in PBS (3 × 5 min), placed in mounting media, and covered with a cover slip. Images were captured with a fluorescent microscope (Eclipse Ti-U, Nikon, Tokyo, Japan). All analyzed images contained 334 ± 14 fibers. The ratio of the number of stained fibers to the total fiber number was calculated. Fibers stained in serial sections with antibodies against slow and fast isoforms were considered hybrid fibers. The CSAs of fast- and slow-twitch muscle fibers were evaluated using ImageJ software (NIH, USA).

Muscle fiber composition in 287 Finnish individuals was estimated based on the expression of MYH1, MYH2, MYH7, Ca^2+^ ATPase A1, and Ca^2+^ ATPase A2 genes, as previously described (*67*). Muscle samples were obtained from the vastus lateralis using a conchotome, under local anesthesia with 20 mg·ml^−1^ lidocaine hydrochloride without epinephrine.

### RNA-seq Analysis of Human Muscle Biopsy Samples

Russian participants were asked not to train for one day before the muscle biopsy of vastus lateralis of the left leg to analyze their gene expression profiles at the resting state. The RNeasy Mini Fibrous Tissue Kit (Qiagen) was used to isolate RNA from 24 muscle tissue samples. Frozen tissue samples were placed in a box submerged in liquid nitrogen. Each frozen sample was transferred onto a sterile Petri dish placed on a frozen plastic ice pack. A piece of tissue with a weight of 10 mg was separated with a sterile scalpel and immediately placed in a 2 mL safe-lock microcentrifuge tube containing 300 µL of lysis buffer and one sterile stainless-steel bead with a diameter of 4 mm. Samples were homogenized using the TissueLyser II system (Qiagen) and shaken twice for 2 min at 25 Hz. RNA samples were isolated according to the manufacturer’s guidelines. RNA concentration was measured using the Qubit spectrophotometer (Thermo Fisher Scientific). RNA quality was assessed using the BioAnalyzer electrophoresis system and BioAnalyzer RNA Nano assay (Agilent Technologies, Santa Clara, CA, USA). RNA integrity number (RIN) was calculated for each RNA sample. Only RNA samples with RIN > 7 were included in the study. Samples were stored at −80 °C until sequencing libraries were prepared. Total RNA samples were treated with DNase I using the Turbo DNA-free Kit (Thermo Fisher Scientific) according to the manufacturer’s instructions. Libraries for RNA sequencing were prepared using the Illumina NEBNext Ultra II Directional RNA Library Prep Kit with the NEBNext rRNA Depletion Module (New England Biolabs). RNA libraries were sequenced on the Illumina HiSeq system with 250 cycles. Sequenced reads were pseudoaligned to the hg38 gencode (v37) transcriptome using kallisto v0.48.0 (*68*) with default settings. Gene-level expression abundances were calculated using the tximport Bioconductor package (*69*). Expression of the MAF genes is presented in transcripts per kilobase million (TPM).

RNA isolation and sequencing in the Finnish samples was performed using strand-specific mRNA-seq, as previously described (*67*). Using the basic GENCODE v19 annotations (*70*), we counted fragments mapping to each gene using htseq-count v0.5.4 (*71*) and quantified gene expression as TPM.

### Statistical Analysis

Data are presented as the mean ± SEM, as indicated in the Figure legends. Statistical significance was determined using GraphPad Prism software (v10.4.0) through Student’s *t*-test or one-way analysis of variance with Tukey’s test. *P < 0.05, **P < 0.01, ***P < 0.001; ns, not significant.

## Acknowledgments

We would like to thank Dr. Pascal Maire (Institut Cochin, INSERM, France) and Dr. Stefano Schiaffino (University of Padova, Italy) for their insightful comments on our study. We also thank Ms. Aki Matsumoto (Kyoto University) for technical assistance with cell culture and sorting and Dr. Takashi Matsuzaka (University of Tsukuba) for technical support with seahorse flux analyzer. We thank the FRCC PCM “Genomics, Proteomics, and Metabolomics” Center for their assistance with RNA sequencing. We also thank Dr. Tsuchimoto and Dr. Kodama (Aichi Medical University, Institute of comprehensive medical research, Division of advanced research promotion) for their assistance in Zeno SWATH-MS data acquisition.

## Author contributions

Methodology: SS, RF

Investigation: SS, RT, TH, RI, MW, GW, IS, KO, TK, HS, MM, EAS, RIS, AVZ, EVG, IIA, RF

Writing—original draft: SS, RF

Funding acquisition: SS, KO, IIA, HS, ST, RF

Conceptualization: SS, RF

All authors read and approved the final manuscript.

## Data and materials availability

All unique reagents generated in this study are available from the corresponding author, Ryo Fujita, in accordance with the relevant material transfer agreements.

- RNA-seq data of human iMuSC-derived myotubes are available in the NCBI Gene Expression Omnibus (GEO) database under accession number “GSE283853”. RNA-seq data of human biopsy are available in the NCBI Gene Expression Omnibus (GEO) database and the Database of Genotypes and Phenotypes under accession numbers “GSE200398” and “phs001048.v2.p1”, respectively.
- LC-MS/MS data have been deposited to the Proteome Xchange Consortium via jPOSTrepo with the dataset identifier JPST003487.
- This paper does not report original code.
- Any additional information required to reanalyze the data reported in this paper is available from the corresponding author, Ryo Fujita upon request.

## References

1. S. Ciciliot, A. C. Rossi, K. A. Dyar, B. Blaauw, S. Schiaffino, Muscle type and fiber type specificity in muscle wasting. Int. J. Biochem. Cell Biol. 45, 2191–2199 (2013).

2. S. Schiaffino, C. Reggiani, Fiber Types in Mammalian Skeletal Muscles. Physiol. Rev. 91, 1447–1531 (2011).

3. R. Bottinelli, M. Canepari, M. A. Pellegrino, C. Reggiani, Force-velocity properties of human skeletal muscle fibres: myosin heavy chain isoform and temperature dependence. J. Physiol. 495, 573–586 (1996).

4. S. Schiaffino, F. Chemello, C. Reggiani, The Diversity of Skeletal Muscle Fiber Types. Cold Spring Harb. Perspect. Biol., a041477 (2024).

5. P. H. Albers, A. J. T. Pedersen, J. B. Birk, D. E. Kristensen, B. F. Vind, O. Baba, J. Nøhr, K. Højlund, J. F. P. Wojtaszewski, Human muscle fiber type-specific insulin signaling: impact of obesity and type 2 diabetes. Diabetes 64, 485–497 (2015).

6. M. Schuler, F. Ali, C. Chambon, D. Duteil, J.-M. Bornert, A. Tardivel, B. Desvergne, W. Wahli, P. Chambon, D. Metzger, PGC1α expression is controlled in skeletal muscles by PPARβ, whose ablation results in fiber-type switching, obesity, and type 2 diabetes. Cell Metab. 4, 407–414 (2006).

7. J. T. Selsby, K. J. Morine, K. Pendrak, E. R. Barton, H. L. Sweeney, Rescue of Dystrophic Skeletal Muscle by PGC-1α Involves a Fast to Slow Fiber Type Shift in the mdx Mouse. PLoS ONE 7, e30063 (2012).

8. N. L. Reyes, G. B. Banks, M. Tsang, D. Margineantu, H. Gu, D. Djukovic, J. Chan, M. Torres, H. D. Liggitt, D. K. Hirenallur-S, D. M. Hockenbery, D. Raftery, B. M. Iritani, Fnip1 regulates skeletal muscle fiber type specification, fatigue resistance, and susceptibility to muscular dystrophy. Proc. Natl. Acad. Sci. 112, 424–429 (2015).

9. T. Hayashi, R. Fujita, R. Okada, M. Hamada, R. Suzuki, S. Fuseya, J. Leckey, M. Kanai, Y. Inoue, S. Sadaki, A. Nakamura, Y. Okamura, C. Abe, H. Morita, T. Aiba, T. Senkoji, M. Shimomura, M. Okada, D. Kamimura, A. Yumoto, M. Muratani, T. Kudo, D. Shiba, S. Takahashi, Lunar gravity prevents skeletal muscle atrophy but not myofiber type shift in mice. Commun. Biol. 6, 424 (2023).

10. R. Jerkovic, C. Argentini, A. Serrano-Sanchez, C. Cordonnier, S. Schiaffino, Early Myosin Switching Induced by Nerve Activity in Regenerating Slow Skeletal Muscle. Cell Struct. Funct. 22, 147–153 (1997).

11. J. Shimizu, F. Kawano, Exercise-induced histone H3 trimethylation at lysine 27 facilitates the adaptation of skeletal muscle to exercise in mice. J. Physiol. 600, 3331–3353 (2022).

12. R. Furrer, B. Heim, S. Schmid, S. Dilbaz, V. Adak, K. J. V. Nordström, D. Ritz, S. A. Steurer, J. Walter, C. Handschin, Molecular control of endurance training adaptation in male mouse skeletal muscle. Nat. Metab. 5, 2020–2035 (2023).

13. J. Lin, H. Wu, P. T. Tarr, C.-Y. Zhang, Z. Wu, O. Boss, L. F. Michael, P. Puigserver, E. Isotani, E. N. Olson, B. B. Lowell, R. Bassel-Duby, B. M. Spiegelman, Transcriptional co-activator PGC-1α drives the formation of slow-twitch muscle fibres. Nature 418, 797–801 (2002).

14. L. Larsson, D. Biral, M. Campione, S. Schiaffino, An age-related type IIB to IIX myosin heavy chain switching in rat skeletal muscle. Acta Physiol. Scand. 147, 227–234 (1993).

15. M. J. Petrany, C. O. Swoboda, C. Sun, K. Chetal, X. Chen, M. T. Weirauch, N. Salomonis, D. P. Millay, Single-nucleus RNA-seq identifies transcriptional heterogeneity in multinucleated skeletal myofibers. Nat. Commun. 11, 6374 (2020).

16. Y. Lai, I. Ramírez-Pardo, J. Isern, J. An, E. Perdiguero, A. L. Serrano, J. Li, E. García-Domínguez, J. Segalés, P. Guo, V. Lukesova, E. Andrés, J. Zuo, Y. Yuan, C. Liu, J. Viña, J. Doménech-Fernández, M. C. Gómez-Cabrera, Y. Song, L. Liu, X. Xu, P. Muñoz-Cánoves, M. A. Esteban, Multimodal cell atlas of the ageing human skeletal muscle. Nature 629, 154–164 (2024).

17. M. Murgia, L. Toniolo, N. Nagaraj, S. Ciciliot, V. Vindigni, S. Schiaffino, C. Reggiani, M. Mann, Single Muscle Fiber Proteomics Reveals Fiber-Type-Specific Features of Human Muscle Aging. Cell Rep. 19, 2396–2409 (2017).

18. L. Wang, P. Gao, C. Li, Q. Liu, Z. Yao, Y. Li, X. Zhang, J. Sun, C. Simintiras, M. Welborn, K. McMillin, S. Oprescu, S. Kuang, X. Fu, A single-cell atlas of bovine skeletal muscle reveals mechanisms regulating intramuscular adipogenesis and fibrogenesis. J. Cachexia Sarcopenia Muscle 14, 2152–2167 (2023).

19. L. Wang, Y. Zhou, Y. Wang, T. Shan, Integrative cross-species analysis reveals conserved and unique signatures in fatty skeletal muscles. Sci. Data 11, 290 (2024).

20. M. A. Pellegrino, M. Canepari, R. Rossi, G. D’Antona, C. Reggiani, R. Bottinelli, Orthologous myosin isoforms and scaling of shortening velocity with body size in mouse, rat, rabbit and human muscles. J. Physiol. 546, 677–689 (2003).

21. V. Smerdu, I. Karsch-Mizrachi, M. Campione, L. Leinwand, S. Schiaffino, Type IIx myosin heavy chain transcripts are expressed in type IIb fibers of human skeletal muscle. Am. J. Physiol.-Cell Physiol. 267, C1723–C1728 (1994).

22. B. Stirn Kranjc, V. Smerdu, I. Eržen, Histochemical and immunohistochemical profile of human and rat ocular medial rectus muscles. Graefes Arch. Clin. Exp. Ophthalmol. 247, 1505–1515 (2009).

23. M. J. Horton, C. Rosen, J. M. Close, J. J. Sciote, Quantification of Myosin Heavy Chain RNA in Human Laryngeal Muscles: Differential Expression in the Vertical and Horizontal Posterior Cricoarytenoid and Thyroarytenoid. The Laryngoscope 118, 472–477 (2008).

24. D. Bloemberg, J. Quadrilatero, Rapid Determination of Myosin Heavy Chain Expression in Rat, Mouse, and Human Skeletal Muscle Using Multicolor Immunofluorescence Analysis. PLOS ONE 7, e35273 (2012).

25. M. Dos Santos, S. Backer, F. Auradé, M. M.-K. Wong, M. Wurmser, R. Pierre, F. Langa, M. Do Cruzeiro, A. Schmitt, J.-P. Concordet, A. Sotiropoulos, F. Jeffrey Dilworth, D. Noordermeer, F. Relaix, I. Sakakibara, P. Maire, A fast Myosin super enhancer dictates muscle fiber phenotype through competitive interactions with Myosin genes. Nat. Commun. 13, 1039 (2022).

26. S. Sadaki, R. Fujita, T. Hayashi, A. Nakamura, Y. Okamura, S. Fuseya, M. Hamada, E. Warabi, A. Kuno, A. Ishii, M. Muratani, R. Okada, D. Shiba, T. Kudo, S. Takeda, S. Takahashi, Large Maf transcription factor family is a major regulator of fast type IIb myofiber determination. Cell Rep. 42, 112289 (2023).

27. F. B. Bastian, J. Roux, A. Niknejad, A. Comte, S. S. Fonseca Costa, T. M. de Farias, S. Moretti, G. Parmentier, V. R. de Laval, M. Rosikiewicz, J. Wollbrett, A. Echchiki, A. Escoriza, W. H. Gharib, M. Gonzales-Porta, Y. Jarosz, B. Laurenczy, P. Moret, E. Person, P. Roelli, K. Sanjeev, M. Seppey, M. Robinson-Rechavi, The Bgee suite: integrated curated expression atlas and comparative transcriptomics in animals. Nucleic Acids Res. 49, D831–D847 (2021).

28. M. Nalbandian, M. Zhao, M. Sasaki-Honda, T. Jonouchi, A. Lucena-Cacace, T. Mizusawa, M. Yasuda, Y. Yoshida, A. Hotta, H. Sakurai, Characterization of hiPSC-Derived Muscle Progenitors Reveals Distinctive Markers for Myogenic Cell Purification Toward Cell Therapy. Stem Cell Rep. 16, 883–898 (2021).

29. M. Zhao, A. Tazumi, S. Takayama, N. Takenaka-Ninagawa, M. Nalbandian, M. Nagai, Y. Nakamura, M. Nakasa, A. Watanabe, M. Ikeya, A. Hotta, Y. Ito, T. Sato, H. Sakurai, Induced Fetal Human Muscle Stem Cells with High Therapeutic Potential in a Mouse Muscular Dystrophy Model. Stem Cell Rep. 15, 80–94 (2020).

30. M. J. Horton, C. A. Brandon, T. J. Morris, T. W. Braun, K. M. Yaw, J. J. Sciote, Abundant expression of myosin heavy-chain IIB RNA in a subset of human masseter muscle fibres. Arch. Oral Biol. 46, 1039–1050 (2001).

31. L. Minati, C. Firrito, A. Del Piano, A. Peretti, S. Sidoli, D. Peroni, R. Belli, F. Gandolfi, A. Romanel, P. Bernabo, J. Zasso, A. Quattrone, G. Guella, F. Lauria, G. Viero, M. Clamer, One-shot analysis of translated mammalian lncRNAs with AHARIBO. eLife 10, e59303 (2021).

32. R. Grifone, C. Laclef, F. Spitz, S. Lopez, J. Demignon, J.-E. Guidotti, K. Kawakami, P.-X. Xu, R. Kelly, B. J. Petrof, D. Daegelen, J.-P. Concordet, P. Maire, Six1 and Eya1 Expression Can Reprogram Adult Muscle from the Slow-Twitch Phenotype into the Fast-Twitch Phenotype. Mol. Cell. Biol. 24, 6253–6267 (2004).

33. C. Niro, J. Demignon, S. Vincent, Y. Liu, J. Giordani, N. Sgarioto, M. Favier, I. Guillet-Deniau, A. Blais, P. Maire, Six1 and Six4 gene expression is necessary to activate the fast-type muscle gene program in the mouse primary myotome. Dev. Biol. 338, 168–182 (2010).

34. K. L. Hetzler, B. C. Collins, R. A. Shanely, H. Sue, M. C. Kostek, The homoeobox gene *SIX1* alters myosin heavy chain isoform expression in mouse skeletal muscle. Acta Physiol. 210, 415–428 (2014).

35. K. Y. Lee, M. K. Singh, S. Ussar, P. Wetzel, M. F. Hirshman, L. J. Goodyear, A. Kispert, C. R. Kahn, Tbx15 controls skeletal muscle fibre-type determination and muscle metabolism. Nat. Commun. 6, 8054 (2015).

36. Z.-X. Meng, S. Li, L. Wang, H. J. Ko, Y. Lee, D. Y. Jung, M. Okutsu, Z. Yan, J. K. Kim, J. D. Lin, Baf60c drives glycolytic metabolism in the muscle and improves systemic glucose homeostasis through Deptor-mediated Akt activation. Nat. Med. 19, 640–645 (2013).

37. Y. Nabeshima, K. Hanaoka, M. Hayasaka, E. Esuml, S. Li, I. Nonaka, Y. Nabeshima, Myogenin gene disruption results in perinatal lethality because of severe muscle defect. Nature 364, 532–535 (1993).

38. M. Dos Santos, A. M. Shah, Y. Zhang, S. Bezprozvannaya, K. Chen, L. Xu, W. Lin, J. R. McAnally, R. Bassel-Duby, N. Liu, E. N. Olson, Opposing gene regulatory programs governing myofiber development and maturation revealed at single nucleus resolution. Nat. Commun. 14, 4333 (2023).

39. J. C. Correia, Y. Kelahmetoglu, P. R. Jannig, C. Schweingruber, D. Shvaikovskaya, L. Zhengye, I. Cervenka, N. Khan, M. Stec, M. Oliveira, J. Nijssen, V. Martínez-Redondo, S. Ducommun, M. Azzolini, J. T. Lanner, S. Kleiner, E. Hedlund, J. L. Ruas, Muscle-secreted neurturin couples myofiber oxidative metabolism and slow motor neuron identity. Cell Metab. 33, 2215–2230.e8 (2021).

40. Y. Deng, L. Lu, H. Zhang, Y. Fu, T. Liu, Y. Chen, The role and regulation of Maf proteins in cancer. Biomark. Res. 11, 17 (2023).

41. K. Huang, M. S. Serria, H. Nakabayashi, S. Nishi, M. Sakai, Molecular cloning and functional characterization of the mouse mafB gene. Gene 242, 419–426 (2000).

42. M. S. Serria, H. Ikeda, K. Omoteyama, J. Hirokawa, S. Nishi, M. Sakai, Regulation and differential expression of the c-maf gene in differentiating cultured cells. Biochem. Biophys. Res. Commun. 310, 318–326 (2003).

43. J.-H. Lee, K. M. Lewis, T. W. Moural, B. Kirilenko, B. Borgonovo, G. Prange, M. Koessl, S. Huggenberger, C. Kang, M. Hiller, Molecular parallelism in fast-twitch muscle proteins in echolocating mammals. Sci. Adv. 4, eaat9660 (2018).

44. J. Janiszewska, M. Bodnar, J. Paczkowska, A. Ustaszewski, M. J. Smialek, L. Szylberg, A. Marszalek, K. Kiwerska, R. Grenman, K. Szyfter, M. Wierzbicka, M. Giefing, M. Jarmuz-Szymczak, Loss of the MAF Transcription Factor in Laryngeal Squamous Cell Carcinoma. Biomolecules 11, 1035 (2021).

45. Y. Na, A. Hall, K. Choi, L. Hu, J. Rose, R. A. Coover, A. Miller, R. F. Hennigan, E. Dombi, M.-O. Kim, S. Subramanian, N. Ratner, J. Wu, MicroRNA-155 contributes to plexiform neurofibroma growth downstream of MEK. Oncogene 40, 951–963 (2021).

46. S. V. Yelamanchili, B. Morsey, E. B. Harrison, D. A. Rennard, K. Emanuel, I. Thapa, D. R. Bastola, H. S. Fox, The evolutionary young miR-1290 favors mitotic exit and differentiation of human neural progenitors through altering the cell cycle proteins. Cell Death Dis. 5, e982–e982 (2014).

47. B. C. Harrison, D. L. Allen, L. A. Leinwand, IIb or not IIb? Regulation of myosin heavy chain gene expression in mice and men. Skelet. Muscle 1, 5 (2011).

48. D. M. Brown, J. M. Brameld, T. Parr, Expression of the Myosin Heavy Chain IIB Gene in Porcine Skeletal Muscle: The Role of the CArG-Box Promoter Response Element. PLOS ONE 9, e114365 (2014).

49. I. Sakakibara, M. Santolini, A. Ferry, V. Hakim, P. Maire, Six Homeoproteins and a linc-RNA at the Fast MYH Locus Lock Fast Myofiber Terminal Phenotype. PLoS Genet. 10, e1004386 (2014).

50. E. R. Chin, E. N. Olson, J. A. Richardson, Q. Yang, C. Humphries, J. M. Shelton, H. Wu, W. Zhu, R. Bassel-Duby, R. S. Williams, A calcineurin-dependent transcriptional pathway controls skeletal muscle fiber type. Genes Dev. 12, 2499–2509 (1998).

51. L. Liu, C. Ding, T. Fu, Z. Feng, J.-E. Lee, L. Xiao, Z. Xu, Y. Yin, Q. Guo, Z. Sun, W. Sun, Y. Mao, L. Yang, Z. Zhou, D. Zhou, L. Xu, Z. Zhu, Y. Qiu, K. Ge, Z. Gan, Histone methyltransferase MLL4 controls myofiber identity and muscle performance through MEF2 interaction. J. Clin. Invest. 130, 4710–4725 (2020).

52. M. J. Potthoff, H. Wu, M. A. Arnold, J. M. Shelton, J. Backs, J. McAnally, J. A. Richardson, R. Bassel-Duby, E. N. Olson, Histone deacetylase degradation andMEF2 activation promote the formation of slow-twitch myofibers. J. Clin. Invest. 117, 2459–2467 (2007).

53. H. Wu, MEF2 responds to multiple calcium-regulated signals in the control of skeletal muscle fiber type. EMBO J. 19, 1963–1973 (2000).

54. S. Schiaffino, Fibre types in skeletal muscle: a personal account. Acta Physiol. 199, 451–463 (2010).

55. K. Okita, Y. Matsumura, Y. Sato, A. Okada, A. Morizane, S. Okamoto, H. Hong, M. Nakagawa, K. Tanabe, K. Tezuka, T. Shibata, T. Kunisada, M. Takahashi, J. Takahashi, H. Saji, S. Yamanaka, A more efficient method to generate integration-free human iPS cells. Nat. Methods 8, 409–412 (2011).

56. M. Zhao, Y. Taniguchi, C. Shimono, T. Jonouchi, Y. Cheng, Y. Shimizu, M. Nalbandian, T. Yamamoto, M. Nakagawa, K. Sekiguchi, H. Sakurai, Heparan Sulfate Chain-Conjugated Laminin-E8 Fragments Advance Paraxial Mesodermal Differentiation Followed by High Myogenic Induction from hiPSCs. Adv. Sci. 11, 2308306 (2024).

57. R. Kawada, T. Jonouchi, A. Kagita, M. Sato, A. Hotta, H. Sakurai, Establishment of quantitative and consistent in vitro skeletal muscle pathological models of myotonic dystrophy type 1 using patient-derived iPSCs. Sci. Rep. 13, 94 (2023).

58. R. Fujita, S. Mizuno, T. Sadahiro, T. Hayashi, T. Sugasawa, F. Sugiyama, Y. Ono, S. Takahashi, M. Ieda, Generation of a MyoD knock-in reporter mouse line to study muscle stem cell dynamics and heterogeneity. iScience 26, 106592 (2023).

59. T. Hayashi, S. Sadaki, R. Tsuji, R. Okada, S. Fuseya, M. Kanai, A. Nakamura, Y. Okamura, M. Muratani, G. Wenchao, T. Sugasawa, S. Mizuno, E. Warabi, T. Kudo, S. Takahashi, R. Fujita, Dual-specificity phosphatases 13 and 27 as key switches in muscle stem cell transition from proliferation to differentiation. Stem Cells Dayt. Ohio, sxae045 (2024).

60. S. Muroya, I. Nakajima, K. Chikuni, Sequential expression of myogenic regulatory factors in bovine skeletal muscle and the satellite cell culture. Anim. Sci. J. 73, 375–381 (2002).

61. K. Ojima, A. Uezumi, H. Miyoshi, S. Masuda, Y. Morita, A. Fukase, A. Hattori, H. Nakauchi, Y. Miyagoe-Suzuki, S. Takeda, Mac-1low early myeloid cells in the bone marrow-derived SP fraction migrate into injured skeletal muscle and participate in muscle regeneration. Biochem. Biophys. Res. Commun. 321, 1050–1061 (2004).

62. D. W. Huang, B. T. Sherman, R. A. Lempicki, Systematic and integrative analysis of large gene lists using DAVID bioinformatics resources. Nat. Protoc. 4, 44–57 (2009).

63. J. Alquicira-Hernandez, J. E. Powell, *Nebulosa* recovers single-cell gene expression signals by kernel density estimation. Bioinformatics 37, 2485–2487 (2021).

64. V. Demichev, C. B. Messner, S. I. Vernardis, K. S. Lilley, M. Ralser, DIA-NN: neural networks and interference correction enable deep proteome coverage in high throughput. Nat. Methods 17, 41–44 (2020).

65. S. Okuda, Y. Watanabe, Y. Moriya, S. Kawano, T. Yamamoto, M. Matsumoto, T. Takami, D. Kobayashi, N. Araki, A. C. Yoshizawa, T. Tabata, N. Sugiyama, S. Goto, Y. Ishihama, jPOSTrepo: an international standard data repository for proteomes. Nucleic Acids Res. 45, D1107–D1111 (2017).

66. C. E. Grant, T. L. Bailey, W. S. Noble, FIMO: scanning for occurrences of a given motif. Bioinformatics 27, 1017–1018 (2011).

67. D. L. Taylor, A. U. Jackson, N. Narisu, G. Hemani, M. R. Erdos, P. S. Chines, A. Swift, J. Idol, J. P. Didion, R. P. Welch, L. Kinnunen, J. Saramies, T. A. Lakka, M. Laakso, J. Tuomilehto, S. C. J. Parker, H. A. Koistinen, G. Davey Smith, M. Boehnke, L. J. Scott, E. Birney, F. S. Collins, Integrative analysis of gene expression, DNA methylation, physiological traits, and genetic variation in human skeletal muscle. Proc. Natl. Acad. Sci. 116, 10883–10888 (2019).

68. N. L. Bray, H. Pimentel, P. Melsted, L. Pachter, Near-optimal probabilistic RNA-seq quantification. Nat. Biotechnol. 34, 525–527 (2016).

69. C. Soneson, M. I. Love, M. D. Robinson, Differential analyses for RNA-seq: transcript-level estimates improve gene-level inferences. F1000Research 4, 1521 (2016).

70. J. Harrow, A. Frankish, J. M. Gonzalez, E. Tapanari, M. Diekhans, F. Kokocinski, B. L. Aken, D. Barrell, A. Zadissa, S. Searle, I. Barnes, A. Bignell, V. Boychenko, T. Hunt, M. Kay, G. Mukherjee, J. Rajan, G. Despacio-Reyes, G. Saunders, C. Steward, R. Harte, M. Lin, C. Howald, A. Tanzer, T. Derrien, J. Chrast, N. Walters, S. Balasubramanian, B. Pei, M. Tress, J. M. Rodriguez, I. Ezkurdia, J. Van Baren, M. Brent, D. Haussler, M. Kellis, A. Valencia, A. Reymond, M. Gerstein, R. Guigó, T. J. Hubbard, GENCODE: The reference human genome annotation for The ENCODE Project. Genome Res. 22, 1760–1774 (2012).

71. S. Anders, P. T. Pyl, W. Huber, HTSeq—a Python framework to work with high-throughput sequencing data. Bioinformatics 31, 166–169 (2015).

