## Supplementary material for "Large MAF Transcription Factors Reawaken Evolutionarily Dormant Fast-Glycolytic Type IIb Myofibers in Human Skeletal Muscle": Fig.S

Shunya Sadaki *et al*.

*Corresponding author. Ryo Fujita


**This PDF file includes:**

Figs. S1 to S5
Tables S1

**Supplementary Figures**

**Fig. S1.**

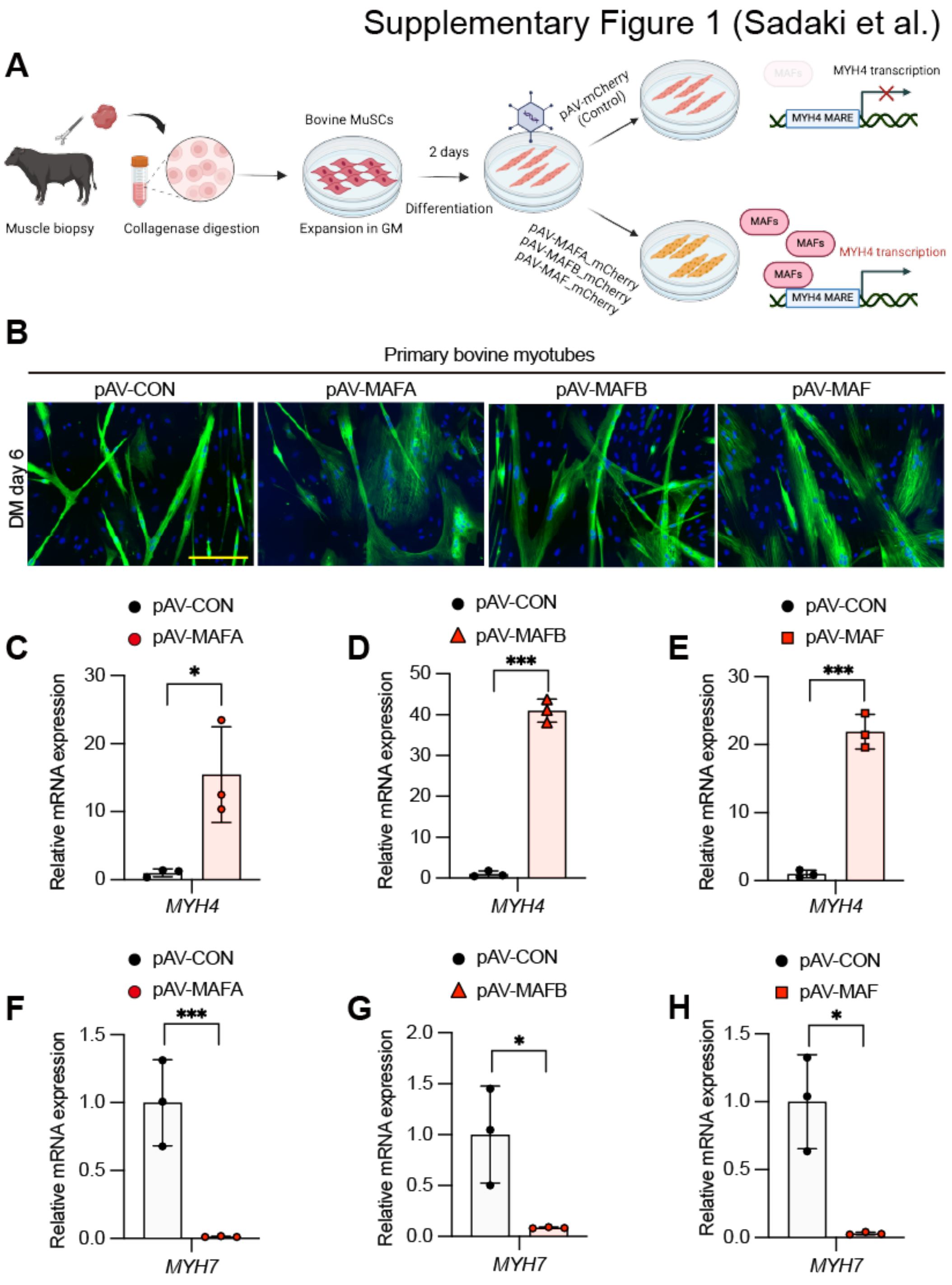


**Fig. S1. Large MAFs can specifically induce MYH4 mRNA expression in bovine MuSCs.** (**A**) A schematic diagram illustrating adenovirus-mediated overexpression of MAFA, MAFB, and MAF in bovine myotubes. Bovine MuSCs were isolated from the thoracic longissimus muscle and cultured in growth medium (GM) for expansion. After two days of exposure to

differentiation medium (DM), bovine myotubes were treated with adenoviral particles expressing MAFA_mCherry (pAV-MAFA), MAFB_mCherry (pAV-MAFB), or MAF_mCherry (pAV-MAF). mCherry-only was used as the control (pAV-CON). Created in BioRender. Ryo, F. (2024) https://BioRender.com/z37l855. (**B**) Immunostaining of bovine MuSCs transduced with pAV-CON, pAV-MAFA, pAV-MAFB, or pAV-MAF, using antibodies against alpha-actinin (α-ACTININ, green). Nuclei were stained with DAPI (blue). (**C-E)** Relative mRNA expression levels of *MYH4*, as determined using RT-qPCR in bovine myotubes four days after transduction with adenoviral vectors expressing MAFA (**C**), MAFB (**D**), or MAF (**E**). mCherry-only was used as the control (pAV-CON, n = 3/group). (**F-H**) Relative mRNA expression levels of *MYH7* in bovine myotubes, as determined using RT-qPCR four days after transduction with adenoviral vectors expressing MAFA (**F**), MAFB (**G**), or MAF (**H**). mCherry-only was used as the control (pAV-CON, n = 3/group). All data are expressed as the mean ± standard error of the mean (SE)
P values were calculated using Student's t-test (**C-H**); *P < 0.05, ***P < 0.001.

**Fig. S2.**

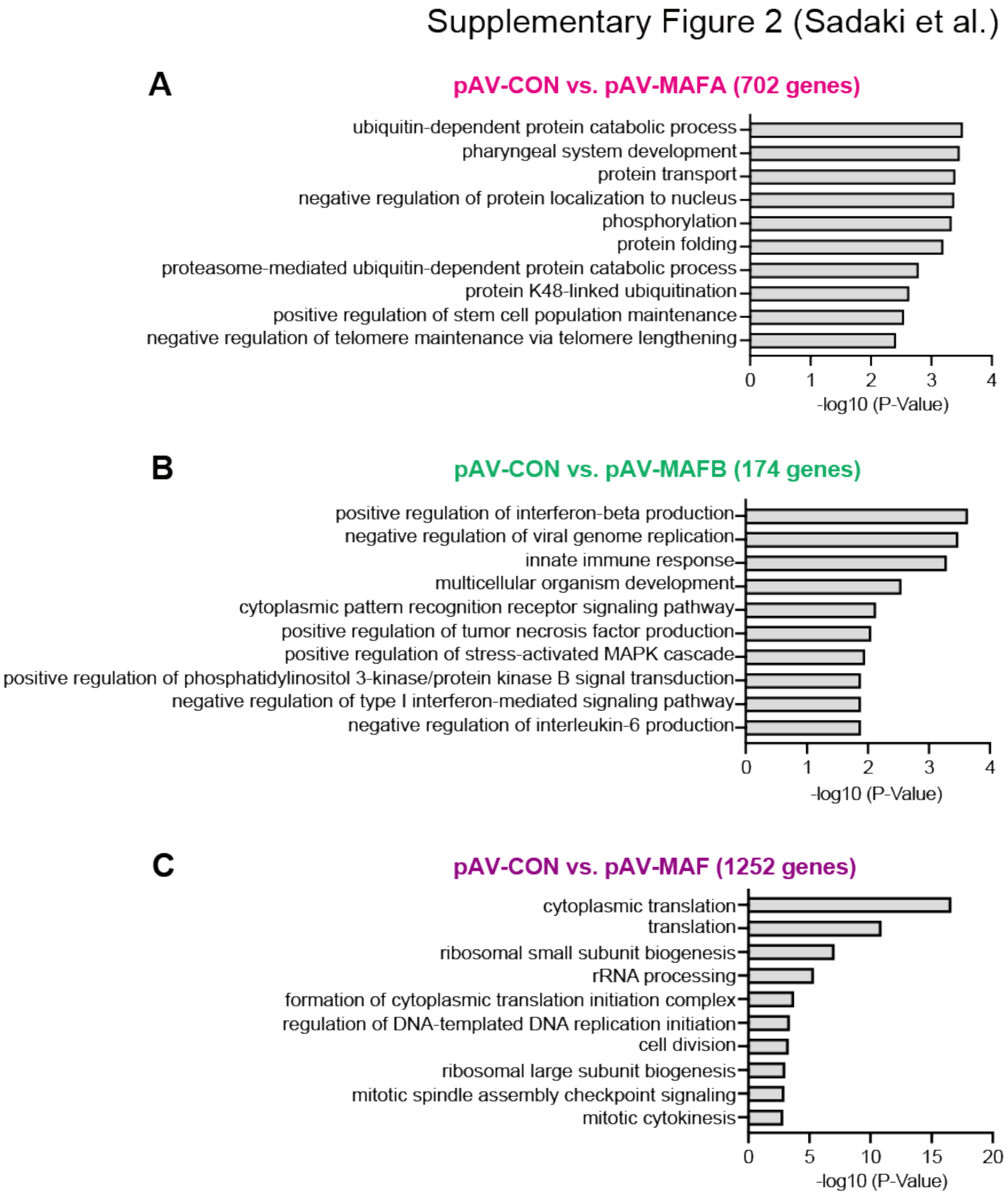


**Fig. S2. Distinct gene sets specifically regulated by MAFA, MAFB, and MAF in human myotubes.** (**A**) Gene Ontology (GO) analysis of 702 MAFA-specific genes identified in Figure 3A. (**B**) GO analysis of 174 MAFB-specific genes identified in Figure 3A. (**C**) GO analysis of 1252 MAF-specific genes identified in Figure 3A.

**Fig. S3.**

Supplementary Figure 3 (Sadaki et al.)

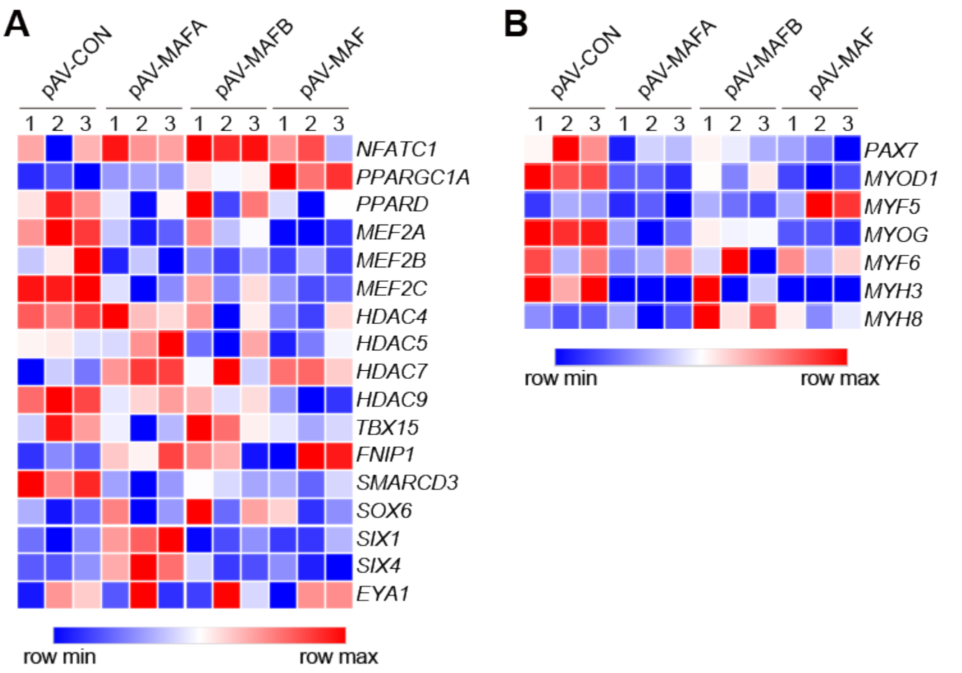


**Fig. S3. Most genes related to fiber type transition and myogenesis were not induced by large MAFs in human myotubes.** (**A**) A heatmap visualizing the expression of myofiber type-regulating genes (n = 3 / group). (**B**) A heatmap visualizing the expression of genes involved in myogenesis (n = 3 / group).

**Fig. S4.**

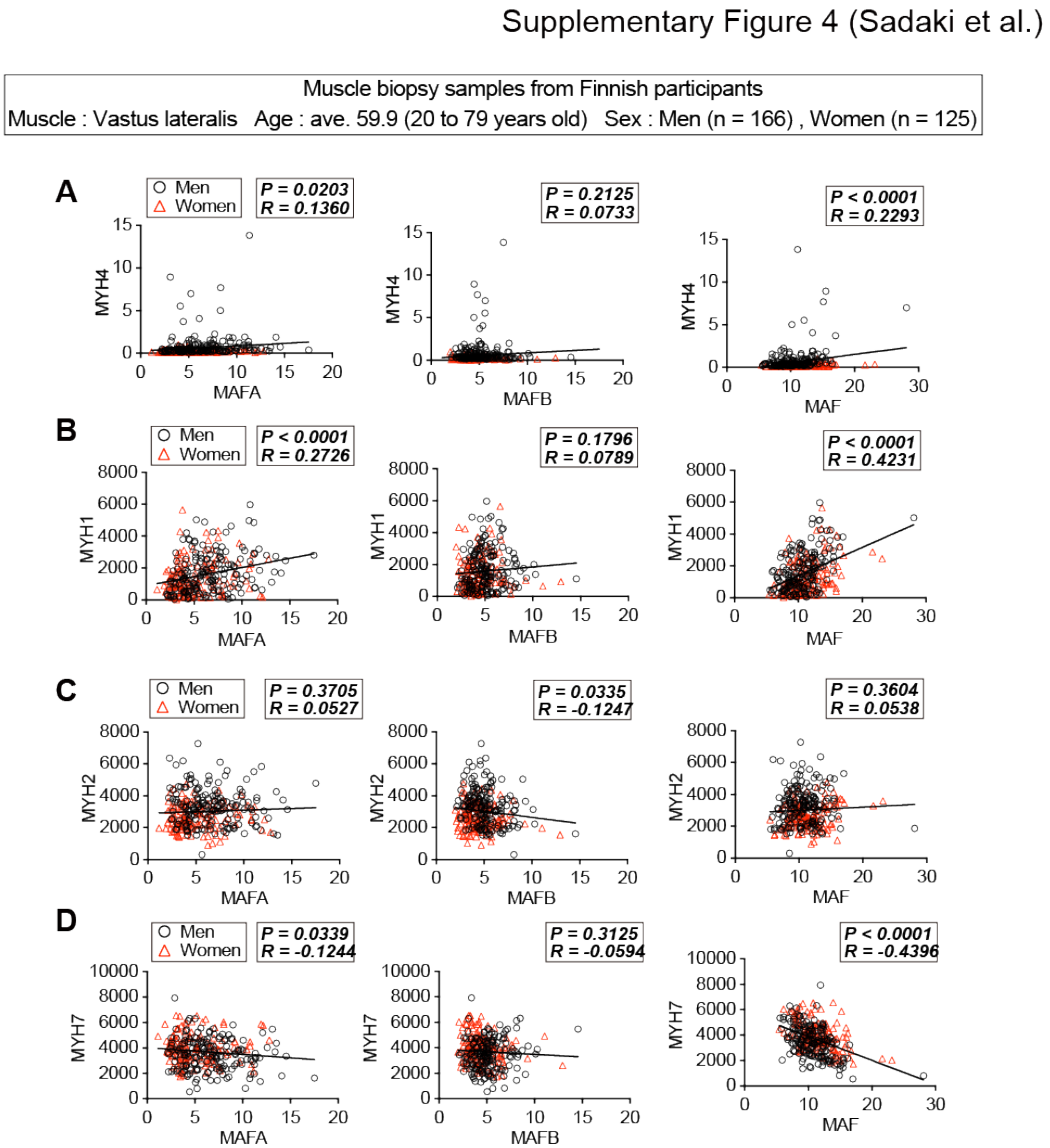


**Fig. S4. Positive correlation of MAFA and MAF expression with MYH4 and MYH1 in muscle biopsy samples of Finnish participants.** Scatter plots showing the mRNA expression levels of *MYH4* (**A**), *MYH1* (**B**), *MYH2* (**C**), and *MYH7* (**D**) (y-axes) versus the combined mRNA expression levels of *MAFA*, *MAFB*, and *MAF* (x-axis) across 291 participants. Pearson correlation coefficients (R) and two-sided P-values (P) were calculated for all analyses (**A**–**D**).

Fig. S5.

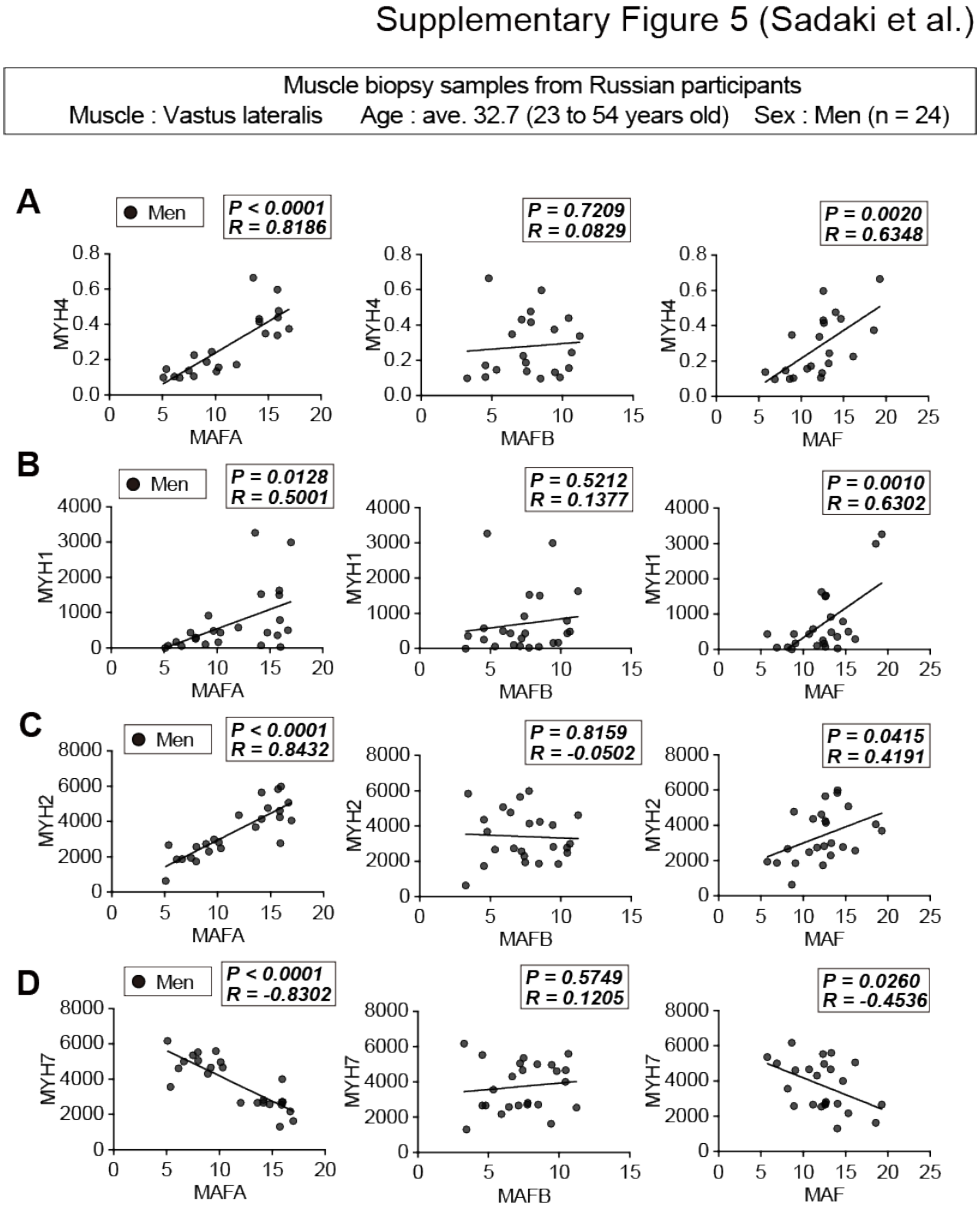


**Fig. S5. Positive correlation of MAFA and MAF expression with MYH4, MYH1, and MYH2 in muscle biopsy samples of Russian participants.** (**A**) Scatter plots showing the mRNA expression levels of *MYH4* (y-axis) versus the combined mRNA expression levels of *MAFA*, *MAFB*, and *MAF* (x-axis) across 21 participants. Outliers were identified using Grubbs' test (Alpha = 0.05); three subjects with outlying MYH4 expression levels were excluded from the analysis. Scatter plots showing the mRNA expression levels of *MYH1* (**B**), *MYH2* (**C**), *MYH7* (**D**) (y-axes) versus the combined mRNA expression levels of *MAFA*, *MAFB*, and *MAF* (x-axis) across 24 participants. Pearson correlation coefficients (R) and two-sided P-values (P) were calculated for all analyses (**A–D**).

**Table S1. List of oligonucleotide sequences used as primers**

| Gene | Fw | Rv |
|---|---|---|
| *MAFA* | TTCAGCAAGGAGGAGGTCAT | AGTTGGCACTTCTCGCTCTC |
| *MAFB* | GTATGTCAACGACTTCGACCTG | ATCCTCGAGGTGTGTCTTCTGT |
| *MAF* | CCTGGCCATGGAATATGTTAAT | AGCCGGTCATCCAGTAGTAGTC |
| *MYH4* | AAAGGTGGCCATTTACAAGCT | CAGCAGAGTTCAGACTTGTCAG |
| *MYH1* | CCCTACAAGTGGTTGCCAGTG | CTTCCCTGCGCCAGATTCTC |
| *MYH2* | CTGGCTGGAGAAGAACAAGG | CAGTTTGAGCCCCAGAGAAG |
| *MYH7* | TGATCTGGAGCTGACACTGG | CTTCTCCTTGGTCAGCTTGG |
| *TBP* | TGTATCCACAGTGAATCTTGGTTG | GGTTCGTGGCTCTCTTATCCTC |
| *MYH7_bovine* | ATAGGCTGGCTGCAGAAAAA | GCTGAGCATCTTGAGGGAAG |
| *MYH4_bovine* | TGACATTGACCACACCCAGT | AGTGCGTGTGATGAGCTGAG |
| *TBP_bovine* | TGAGCCAGAGTTATTTCCTGGT | TCTGCTCTGACTTTAGCACCTG |
